# Hybrid speciation and ghost ancestry shape the *Anopheles gambiae* species complex

**DOI:** 10.64898/2026.08.20.745903

**Authors:** Yang Yang, Xiao-Xu Pang, Wei-Ning Bai, Bo-Wen Zhang, Da-Yong Zhang

## Abstract

Speciation within reticulate radiations can involve both lineage divergence and hybrid lineage formation, yet recurrent introgression obscures both histories. In the *Anopheles gambiae* complex, gene-family presence-absence data yielded a species tree favored over four sequence-derived alternatives by network-model comparison. D-BPP analyses recovered seven reticulation events, including multiple ghost-lineage contributions, and supported a ghost-mediated hybrid origin of *A. merus*. Simulations showed that sampled-parent hybrid origin generates temporal convergence between reticulation and lineage formation when analyzed under an ordinary introgression model; this signature supported hybrid speciation in *A. gambiae*. Loci with contrasting parental affinities contained olfactory and cuticular genes with potential roles in prezygotic isolation. Together, these results resolve species relationships and identify candidate genomic mechanisms through which hybridization may have contributed to reproductive isolation.

## Introduction

Species can arise through fundamentally different evolutionary histories. In a purely bifurcating divergence history, an ancestral lineage splits and its descendants progressively diverge, producing the branching pattern represented by a species tree (Kubatko 2026). In homoploid hybrid speciation (HHS), differentiated parental lineages hybridize and jointly give rise to a new lineage that begins an independent evolutionary trajectory without a change in ploidy. This independence may arise through reproductive barriers generated by hybridization itself (Schumer et al. 2014; Long and Rieseberg 2025) or be initially maintained by geographic or ecological isolation, with reproductive isolation evolving later (Nieto Feliner et al. 2017; Ottenburghs 2018). Distinguishing these histories is difficult in rapid radiations, where recurrent introgression overlays secondary genetic exchange on the original process of lineage formation (Kong et al. 2025; Schumer and Rieseberg 2026; Solís-Lemus 2026). The *Anopheles gambiae* complex exemplifies this problem: rapid diversification, incomplete lineage sorting (ILS), and extensive interspecific gene flow (Fontaine et al. 2015; Wen et al. 2016; Thawornwattana et al. 2018) have left both its species-divergence history and the origins of individual lineages unresolved.

Recovering the species-divergence history is the first challenge. Nucleotide sequences record both lineage divergence and subsequent genetic exchange, so introgressed haplotypes can shift genealogical signal away from the underlying branching history (Mallet et al. 2016; Jiao et al. 2021; Hibbins and Hahn 2022), while rapid radiation further generates extensive ILS. Most empirical studies infer a bifurcating species tree first and then add reticulation edges to that fixed topology (Francis and Steel 2015; Solís-Lemus 2026). Errors in the starting tree can propagate through network inference, while ghost lineages can cause unsampled ancestry to be misattributed to gene flow among sampled taxa (Tricou et al. 2022; Pang and Zhang 2024; Cheng et al. 2026; Yang et al. 2026).

Gene-family presence–absence variation may provide an alternative source of species-tree information because it responds differently to introgression than nucleotide similarity (Fig. 1). Gene-content characters have long been used in microbial and deep eukaryotic phylogenetics (Fitz-Gibbon and House 1999; Snel et al. 1999; Delsuc et al. 2005; Pett et al. 2019; Zhao et al. 2021), but remain little explored in recent reticulate radiations. Introgression can replace alleles or add gene copies without altering whether an already represented family is present. By contrast, a binary presence–absence state changes only when a previously absent family becomes established or an existing family is completely lost (Novick and Doolittle 2020). Because both processes are constrained by purifying selection in the recipient genome, interspecific gene flow is unlikely to alter these states. Moreover, gene-content matrices aggregate information across thousands of families, reducing the influence of a relatively small subset positively selected in a recipient genome. We therefore presume that gene-family content could faithfully recover species branching order under extensive gene flow and test this presumption by comparing reticulate histories built on competing species trees with our recently introduced D-BPP framework (Yang et al. 2026).

**Fig. 1.**
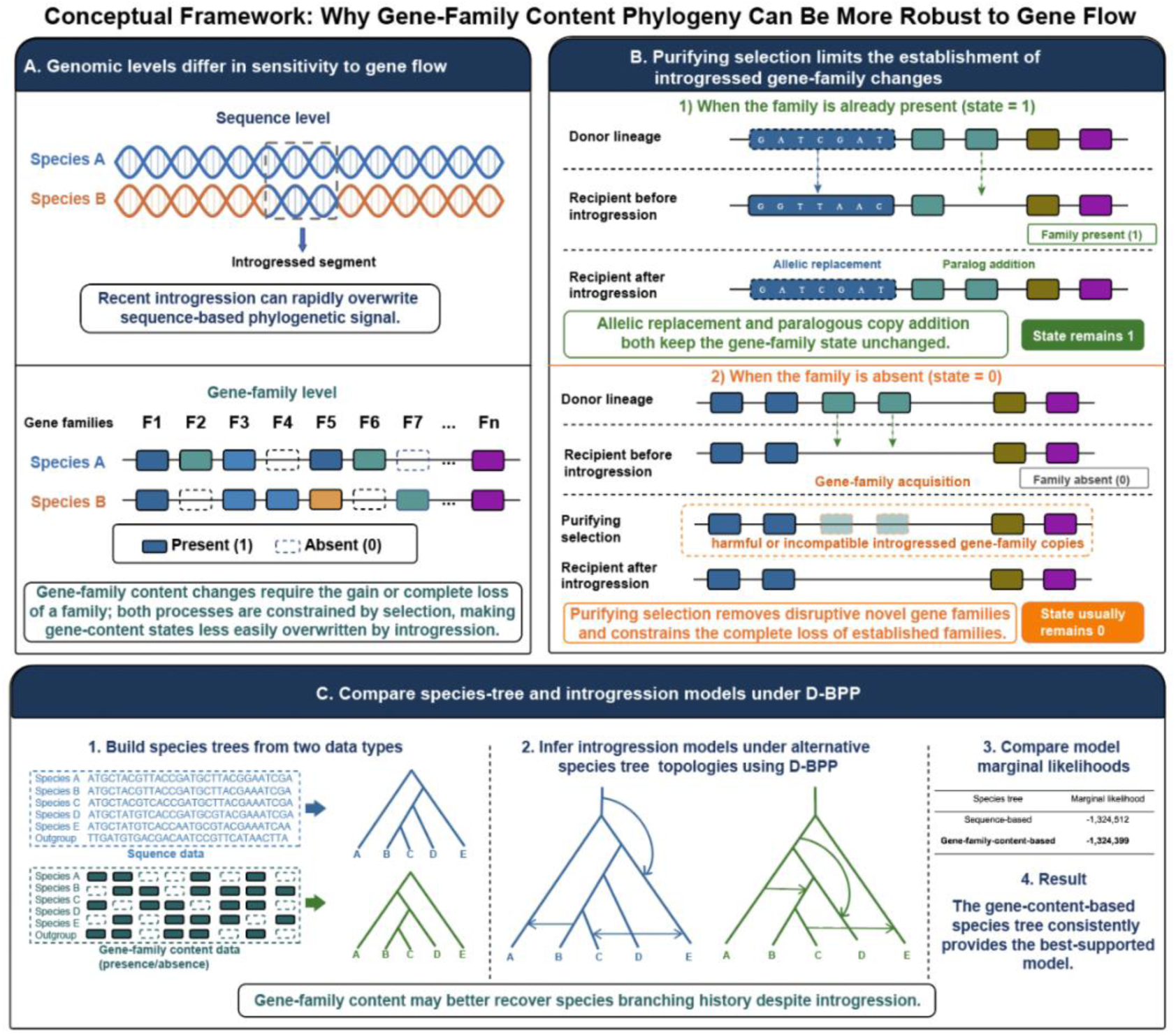
Gene-family content provides an alternative signal of species divergence under introgression. **(A)** Sequence-level and gene-family-level characters differ in their response to introgression. Introgressed genomic segments can alter local sequence genealogies, whereas gene-family content is encoded as binary presence/absence states. Allelic replacement or addition of paralogous copies does not alter the presence state of a family that is already present. Changes in gene-family state require gain and retention of a previously absent family or complete loss of an existing family. **(B)** Selection may constrain the establishment or loss of gene families following introgression. If a family is already present, introgression may replace alleles or add paralogous copies without changing its presence state. If a family is absent, an introgressed family must become established to change the state, whereas complete loss is required to convert a present state to absence. **(C)** D-BPP evaluates reticulate histories built on alternative candidate species trees inferred from sequence data and gene-family content. Marginal-likelihood comparison identifies the topology-network model best supported by the multilocus sequence data.

Identifying hybrid-origin events is a second challenge. Most phylogenetic-network models parameterize lineage divergence and reticulation as separate events, so a hybrid-origin history may be represented as ordinary introgression unless their temporal relationship is examined explicitly. Hybrid origin predicts that hybridization coincides with the origin of the new lineage, whereas secondary introgression occurs after the recipient lineage has already diverged (Folk et al. 2018; Flouri et al. 2020). This relationship provides a testable signature of hybrid lineage formation, but inferred reticulation times can be biased when event order or ghost-lineage placement is mis-specified (Huang et al. 2022). We therefore used simulations to calibrate these effects before applying the temporal diagnostic to empirical networks.

Hybrid lineage formation and establishment of reproductive isolation need not be simultaneous. A hybrid lineage may become evolutionarily independent because hybridization generates reproductive barriers, or because geographic or ecological separation initially limits gene exchange while reproductive isolation evolves later (Schumer et al. 2014; Nieto Feliner et al. 2017; Ottenburghs 2018; Long and Rieseberg 2025). When hybridization contributes to isolation, parental genetic components involved in reproductive barriers may be assorted and retained in the hybrid lineage (Wang et al. 2021). Genomic regions with contrasting parental affinities and functions relevant to reproductive isolation can therefore identify candidate mechanisms linking hybrid ancestry to lineage independence.

Here, we reconstruct species-divergence histories and test hybrid origins in the *A. gambiae* complex. We generate competing branching hypotheses from gene-family and sequence data, calibrate temporal signatures of hybrid origin and model misspecification by simulation, and reconstruct D-BPP networks on five candidate species trees. Marginal-likelihood comparison favors a gene-family history that places *A. quadriannulatus* as the earliest-diverging ingroup, pairs *A. melas* with *A. merus*, and resolves *A. gambiae* and *A. coluzzii* as sisters with *A. arabiensis* branching earlier. The corresponding network reveals recurrent introgression, multiple ghost contributions, a hybrid origin of *A. gambiae*, and even a ghost-mediated hybrid origin of *A. merus*. We used population-genomic scans and locus-specific phylogenies to identify olfactory- and cuticle-associated genomic regions as targets for testing whether parental alleles underlying interspecific reproductive barriers have undergone alternative assortment in *A. gambiae*, potentially serving as independent evidence supporting hybrid speciation.

## Results

### Gene-family content provides a stable candidate divergence history

Gene-family content consistently recovered the same ingroup topology. Publicly available gene annotations from eight *Anopheles* genomes were processed using a common curation pipeline, and OrthoFinder (Emms and Kelly 2019) inferred 11,052 to 12,905 orthogroups across four minimum protein-length thresholds. All four matrices recovered (*A. quadriannulatus*, ((*A. melas*, *A. merus*), (*A. arabiensis*, (*A. gambiae*, *A. coluzzii*)))), and the topology remained unchanged across 30 perturbation replicates with symmetric false-gain and false-loss probabilities from 0.001 to 0.01 (Fig. 2A; fig. S1; table S1).

**Fig. 2.**
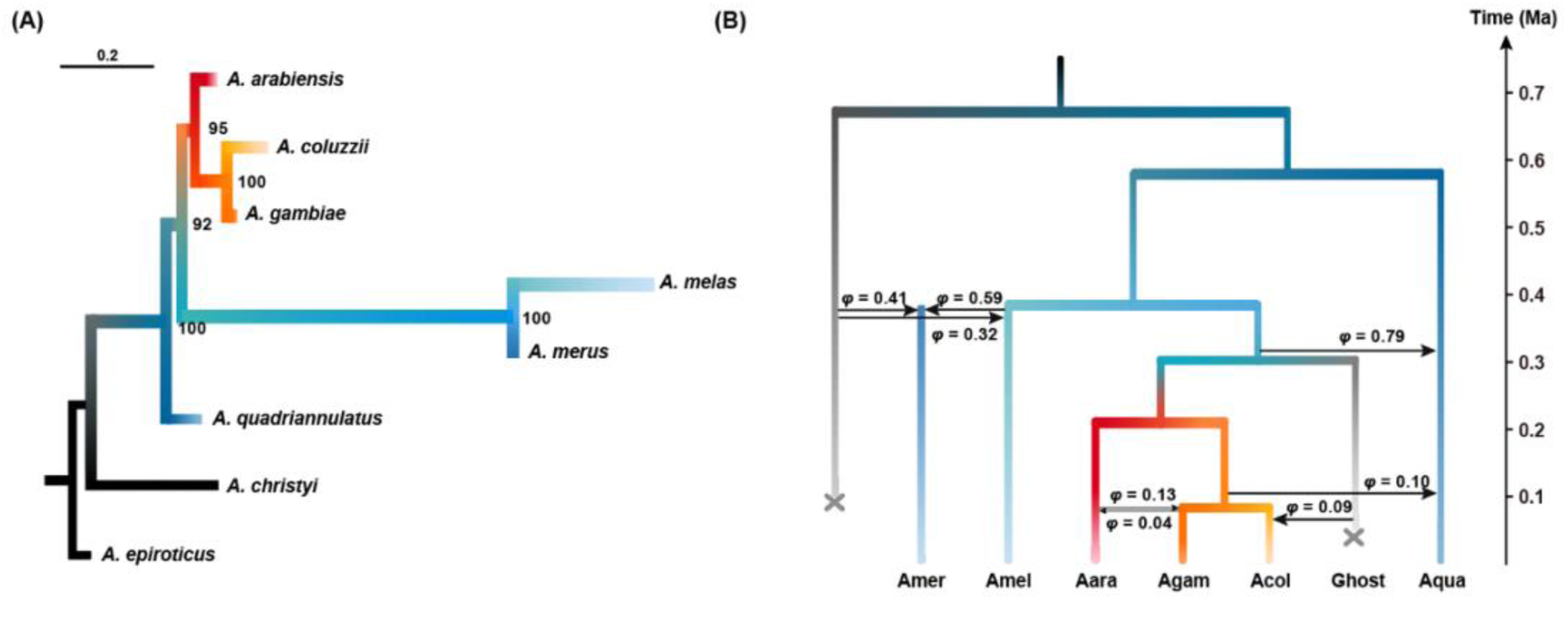
Gene-family species tree and the best-supported reticulate history of the *Anopheles gambiae* complex. **(A)** Gene-family content phylogeny inferred from the orthogroup presence/absence matrix constructed using a minimum protein-length threshold of 100 amino acids. Node labels indicate bootstrap support from 1,000 nonparametric replicates. **(B)** Best-supported reticulate evolutionary network inferred by D-BPP using the gene-family topology as the candidate species tree. Branch lengths are scaled to absolute time in millions of years ago (Ma), using a mutation rate of 2.8 × 10−9 substitutions per site per generation and 11 generations per year (Keightley et al. 2014; Thawornwattana et al. 2018). Gray branches represent inferred unsampled ghost lineages. Arrows indicate retained reticulation events; arrow direction denotes the inferred direction of gene flow, and φ values indicate estimated introgression probabilities or parental contributions. Abbreviations: Amer, *A. merus*; Amel, *A. melas*; Aara, *A. arabiensis*; Acol, *A. coluzzii*; Agam, *A. gambiae*; Aqua, *A. quadriannulatus*.

Sequence data produced alternative branching hypotheses. From 4,214 autosomal, noncoding, unlinked loci, ASTRAL-III (Zhang et al. 2018) recovered a more pectinate topology (fig. S2). We therefore carried forward five candidate species trees: the gene-family topology, the ASTRAL topology, the X-chromosome and whole-genome topologies of Fontaine et al. (2015), and the topology of Thawornwattana et al. (2018). Each candidate tree was evaluated with the same 4,214 loci in the subsequent *D*-statistic, D-BPP, and marginal-likelihood analyses.

### Simulations calibrate reticulation inference and hybrid-origin signatures

Simulation showed that reticulation-model misspecification produces systematic timing biases (Fig. 3). Reversing the order of two introgression events compressed their estimated times toward one another. Assigning ghost introgression to an incorrect recipient branch shifted the inferred reticulation time toward the divergence time of that branch. These patterns provide temporal signatures of incorrect event order and ghost-lineage placement.

**Fig. 3.**
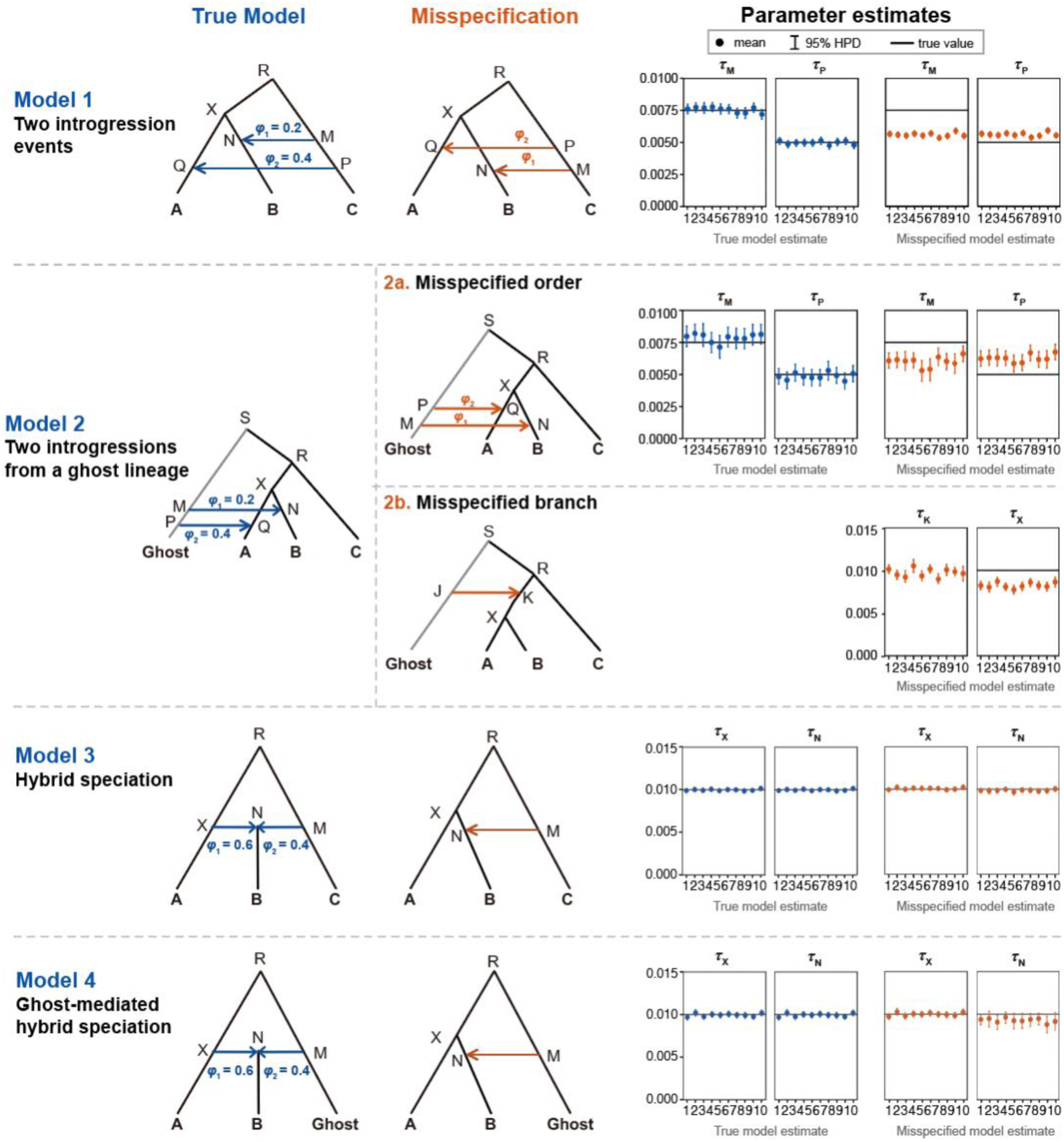
Simulation-calibrated temporal signatures of reticulation-model misspecification and hybrid origin. For each scenario, the generating model is shown on the left, the corresponding misspecified model in the middle, and posterior estimates of key time parameters on the right. Blue arrows and points indicate analyses under the generating model, whereas orange arrows and points indicate analyses under misspecified models. Model 1 represents two introgression events from a sampled donor; Model 2 represents two introgression events from a ghost lineage and evaluates misspecification of event order and recipient branch; Model 3 represents hybrid speciation with both parental lineages sampled; and Model 4 represents ghost-mediated hybrid speciation. Each point indicates the posterior mean from one simulation replicate, vertical bars indicate the corresponding 95% highest posterior density interval, and horizontal black lines indicate the generating parameter values.

Sampled-parent hybrid origin produced a different signature. When hybrid speciation with both parental lineages sampled was analyzed as ordinary introgression, the reticulation and lineage-divergence times converged. Ghost-mediated hybrid speciation produced weaker temporal convergence, making explicit network comparison more informative for ghost-parent cases (Fig. 3).

We calibrated the Savage-Dickey test for H0: τN = τX versus H1: τN < τX, representing temporal equivalence as 0 < τX − τN < ε for ε = 10^−5^ and 10^−6^. Bε was calculated as the ratio of the prior to posterior probability of this interval, with the prior probability estimated from matched BPP runs without sequence data (Thawornwattana et al. 2025). Bε < 0.01 supported temporal equivalence and Bε > 100 supported post-divergence introgression (Thawornwattana et al. 2025). Sampled-parent simulations shifted posterior probability toward temporal coincidence, whereas ghost-mediated simulations were less decisive (table S2). These results defined the empirical inference strategy: sampled-parent candidates were evaluated with temporal tests, whereas ghost-mediated candidates were evaluated by explicit network-model comparison.

### Joint topology-network comparison favors the gene-family species tree

Guided by the simulation-calibrated inference strategy, we reconstructed a complete reticulate history under each of the five candidate species trees. Under the gene-family topology, *D*-statistics identified 13 significant trios (table S3). Bayesian testing retained recurrent gene flow into *A. quadriannulatus*; *A. arabiensis* → *A. gambiae* and *A. gambiae* → *A. arabiensis*; ghost ancestry in *A. coluzzii*; and ghost contributions to *A. merus* and *A. melas* (Fig. 2B; table S4). Screening and reconstruction results for all five starting trees are reported in tables S3 to S8, Supplementary Texts S1 to S5, and fig. S3.

The *A. merus* ghost contribution was inferred close to the *A. merus*-*A. melas* divergence after all retained events were incorporated (table S9). Because the simulations showed that ghost-lineage placement can displace reticulation times toward species divergence, we compared two explicit histories: post-divergence ghost introgression and ghost-mediated hybrid origin. The ghost-mediated hybrid-origin model had the higher marginal likelihood in all four data partitions and was retained in the completed network (table S10).

The completed topology-network models were then compared using marginal likelihoods estimated by thermodynamic integration across the same four nonoverlapping partitions of the 4,214 loci. The network based on the gene-family topology had the highest marginal likelihood in every partition and exceeded the previously published network scenarios evaluated on the same data (fig. S3; table S11). Joint modeling of branching and reticulation therefore favors the gene-family species tree for this radiation.

Posterior estimates from all 4,214 loci placed the sampled-complex crown at ∼0.589 Ma and the ghost-mediated hybrid origin of *A. merus* at ∼0.386 Ma, with estimated parental contributions of 0.59 from the *A. melas* lineage and 0.41 from the ghost lineage. *A. arabiensis* diverged from the *A. gambiae*-*A. coluzzii* ancestor at ∼0.218 Ma, and *A. gambiae* and *A. coluzzii* diverged at ∼0.087 Ma (Fig. 2B; table S12). The completed history contained seven reticulation events. An empirical-scale simulation generated from its posterior mean parameters recovered all seven generating edges through the same D-BPP workflow (tables S13 and S14).

### Temporal coincidence supports a hybrid origin of *A. gambiae*

The timing of *A. arabiensis* ancestry supports a hybrid origin of *A. gambiae*. All five candidate species trees recovered the same local relationship among *A. gambiae*, *A. coluzzii*, and *A. arabiensis*. We therefore analyzed these three species in a reduced network containing *A. arabiensis* → *A. gambiae*, the reverse-direction event, and ghost introgression to *A. coluzzii*, with independent time parameters for the two directional reticulations (Fig. 4A).

**Fig. 4.**
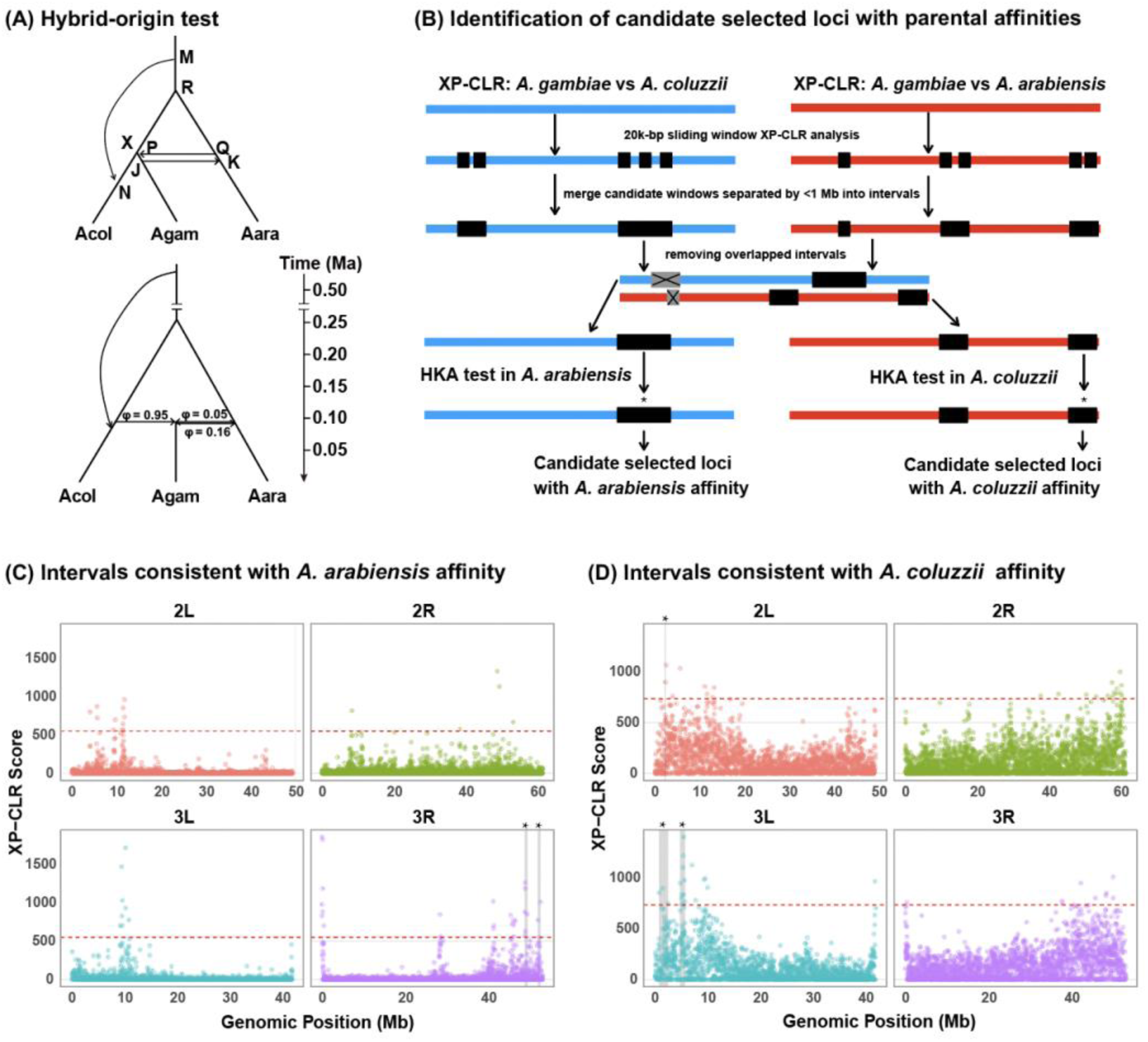
Temporal and genomic evidence for a hybrid origin of *Anopheles gambiae*. **(A)** Reduced three-species model and time-calibrated network for *A. coluzzii* (Acol), *A. gambiae* (Agam), and *A. arabiensis* (Aara). The highlighted reticulation marks the candidate hybridization event; φ values indicate inheritance probabilities. **(B)** Workflow for identifying selected intervals with parental-affinity patterns. Comparison-specific XP-CLR intervals were filtered by HKA tests in the inferred source lineage. **(C, D)** Genome-wide XP-CLR profiles for chromosome arms 2L, 2R, 3L, and 3R. Panel C shows intervals with *A. arabiensis* affinity and panel D intervals with *A. coluzzii* affinity. Red dashed lines mark the upper 0.5% genome-wide XP-CLR threshold; gray shading and asterisks indicate intervals additionally supported by HKA tests.

In the reduced three-species model, the *A. arabiensis* → *A. gambiae* reticulation occurred at ∼0.0912 Ma, nearly coincident with the origin of the *A. gambiae* lineage at the *A. gambiae-A. coluzzii* divergence (∼0.0914 Ma). Savage-Dickey ratios were Bε = 0.0102 for ε = 10^−5^ and Bε = 0.006 for ε = 10^−6^ (table S15). Ten replicate datasets simulated under the empirical three-species network reproduced low Bε values, and none supported post-divergence introgression (table S16). Consistent Bε support for temporal equivalence was also obtained for the *A. arabiensis* → *A. gambiae* event in the full-network analysis (table S12). Together, these results support hybridization coinciding with the origin of the *A. gambiae* lineage rather than representing a later introgression event. The inferred parental contributions were strongly asymmetric, with 0.95 from the lineage leading to *A. coluzzii* and 0.05 from *A. arabiensis* (Fig. 4A).

### Parental-affinity loci identify candidate mechanisms of hybrid isolation

We next asked whether hybridization may also have contributed genetic components involved in maintaining the evolutionary independence of *A. gambiae*. Among the HKA-supported comparison-specific intervals identified by XP-CLR scans followed by HKA validation, we focused on a 3L interval with greater affinity of *A. gambiae* to *A. coluzzii* and a 3R interval with greater affinity to *A. arabiensis*, both of which contained genes potentially relevant to prezygotic reproductive isolation (Fig. 4B to D; table S17).

The 3L interval contained the odorant-binding protein OBP4 (AGAP010489) and odorant receptors Or43 (AGAP010504) and Or44 (AGAP010505) (table S18). Gene trees for OBP4, Or43, and Or44 grouped *A. gambiae* more closely with *A. coluzzii*, matching the regional parental-affinity signal (figs. S4 to S6). OBP4 has experimentally supported odorant-binding activity, and Or43 and Or44 belong to the odorant-receptor family (Davrazou et al. 2011; Qiao et al. 2011). The 3R interval contained the RR-2 cuticular-protein array CPR82-CPR100 (AGAP010095-AGAP010128) (table S18). AGAP010105 provided sufficient phylogenetic information and grouped *A. gambiae* with *A. arabiensis* (fig. S7), matching the regional parental-affinity signal. RR-2 proteins are structural components of rigid arthropod cuticle (Cornman et al. 2008; Togawa et al. 2008; Vannini and Willis 2017). The 3L and 3R regions therefore identify alternate parental genomic components in olfactory and cuticular systems with potential roles in prezygotic isolation. These loci provide explicit targets for testing whether genetic variation brought together through hybridization contributed to the subsequent establishment or maintenance of reproductive isolation in *A. gambiae*.

## Discussion

Homoploid hybrid speciation was long regarded as uncommon (Schumer et al. 2014; Taylor and Larson 2019), but theoretical and empirical work increasingly suggests it may be more widespread than previously recognized (Blanckaert and Bank 2018; Long and Rieseberg 2025; Wang et al. 2025). Documenting HHS requires distinguishing hybrid-lineage formation from secondary introgression and establishing how such lineages attain evolutionary independence. By integrating gene-family-based species-tree inference with D-BPP network reconstruction, simulation-calibrated temporal tests, and population-genomic analyses, we address both challenges within the *Anopheles gambiae* complex. Our reconstructed evolutionary history supports a hybrid origin for *A. gambiae* and a ghost-mediated hybrid origin for *A. merus*. In *A. gambiae*, candidate regions with contrasting parental affinities harbour olfactory and cuticular genes, offering putative genomic links between hybrid ancestry and lineage isolation. Together, these results add to mounting evidence that HHS may be substantially underrecognized (Wang et al. 2025). We note that estimating the true prevalence of HHS remains difficult: hybrid origins must be disentangled from post-divergence introgression, and the mechanisms enabling hybrid lineages to achieve evolutionary independence from their parental species often remain unresolved (Nieto Feliner et al. 2017; Folk et al. 2018; Ottenburghs 2018). These difficulties are further compounded by a historical shortage of analytical frameworks capable of reconstructing complex reticulate histories and formally testing whether hybridization coincided with lineage formation (Folk et al. 2018; Flouri et al. 2020; Thawornwattana et al. 2025).

### Gene-family content provides an alternative species-tree signal for reticulate species complexes

Gene-family content extends a long-established phylogenetic character system from microbes and deeper eukaryotic divergences to a recent radiation dominated by ILS and introgression (Fitz-Gibbon and House 1999; Snel et al. 1999; Delsuc et al. 2005; Pett et al. 2019; Zhao et al. 2021). Its phylogenetic value arises from the different ways in which introgression affects gene-family states and nucleotide haplotypes.

Introgression can immediately replace alleles and reshape local sequence genealogies, whereas changing a binary gene-family state requires establishment of a previously absent family or complete loss of an existing one. Because both the establishment of introgressed gene families and the complete loss of resident families are constrained by selection, interspecific gene flow is less likely to alter presence-absence states than to reshape allele-level variation. Gene gain, loss, duplication, introgression, and annotation error can nevertheless modify presence-absence patterns, making robustness to data processing an important empirical test. In the *A. gambiae* complex, the same topology was recovered across four protein-length thresholds, and throughout perturbation analyses that introduced false gains and losses. The consistency of this topology across these analyses argues against the recovered branching pattern being driven by a particular filtering choice or gene annotation error.

The recovered topology is also consistent with major biological differentiation within the complex. It pairs the coastal brackish-water specialists *A. melas* and *A. merus*, groups *A. gambiae*, *A. coluzzii*, and *A. arabiensis*, and places the largely zoophilic *A. quadriannulatus* as the earliest-diverging ingroup lineage (Coluzzi et al. 1979; White et al. 2011; Wiebe et al. 2017). Sequence-based studies have instead recovered conflicting relationships and have often emphasized the X-chromosome inversion region as a source of species-tree signal less affected by autosomal introgression (Fontaine et al. 2015; Thawornwattana et al. 2018). Low recombination can reduce the effective exchange of introgressed ancestry (Burbrink et al. 2025), but it does not uniquely preserve species-divergence history. Although recombination and introgression are often positively correlated, this relationship can be weak, and low-recombination regions do not necessarily preserve the true species phylogeny because ILS, linked selection, and occasional adaptive introgression can still distort local genealogies (Nater et al. 2015; Hibbins and Hahn 2024). Gene-family content therefore provides an independent genome-wide source of species-tree information rather than relying on a restricted genomic region presumed to retain the branching history.

The strongest support for the gene-family topology came from explicit network-model comparison. Each of the five candidate species trees was used to reconstruct its own D-BPP reticulate history, and the completed topology-network models were evaluated with the same multilocus sequence dataset of 4,214 loci. The gene-family-based network had the highest marginal likelihood in all four data partitions and also exceeded the previously published network scenarios evaluated on the same data. The sequence data therefore independently favored the divergence history recovered from gene-family content once recurrent introgression and ghost ancestry were incorporated into the model. Species-tree inference and network reconstruction can thus be evaluated jointly, and among the alternatives examined here, gene-family content provides the branching history that best reconciles species divergence with the reticulation recorded across the genome.

### Ghost ancestry and hybrid lineage formation shape the radiation

The best-supported network contains seven reticulation events, showing that the diversification of the *A. gambiae* complex involved repeated genetic exchange alongside lineage divergence. Ghost ancestry contributed to *A. merus*, *A. melas*, and *A. coluzzii*, placing extinct or unsampled lineages directly within the evolutionary history of several extant species. The empirical-scale simulation recovered all seven generating edges, showing that this level of reticulation is recoverable with the D-BPP workflow (Yang et al. 2026).

*A. merus* provides the clearest example of ghost ancestry contributing to lineage formation. The ghost contribution was inferred near the origin of the *A. merus* lineage, prompting explicit comparison between post-divergence ghost introgression and ghost-mediated hybrid origin. The ghost-mediated hybrid-origin model was favored in every data partition. The estimated ancestry proportions—0.59 from the *A. melas* lineage and 0.41 from the ghost lineage—indicate substantial contributions from both parental sources. Hybrid lineage formation involving unsampled ancestry therefore contributed to diversification within the complex.

### Temporal coincidence supports a hybrid origin of *A. gambiae*

Hybrid origin with sampled parents leaves a recoverable temporal signature. When sampled-parent hybrid-origin data were analyzed with an ordinary introgression model, the estimated reticulation time converged on the time at which the hybrid lineage originated. Incorrect event order and ghost-branch placement produced distinct timing distortions (Fig. 3), consistent with an earlier finding (Huang et al. 2022). These simulations show that temporal coincidence between reticulation and lineage origin can serve as a diagnostic signature of sampled-parent hybrid formation.

In *A. gambiae*, concordant evidence from the reduced and complete-network analyses supports a hybrid origin rather than secondary introgression into an already established lineage. The temporal coincidence between the *A. arabiensis* contribution and the origin of *A. gambiae* indicates that hybridization accompanied lineage formation. The inferred ancestry was strongly asymmetric, with approximately 5% contributed by *A. arabiensis*, showing that hybrid lineage formation can involve markedly unequal parental contributions. Such asymmetry is also illustrated by a butterfly hybrid-origin system in which one parental lineage contributed >99% of the ancestry and the other <1% (Rosser et al. 2024).

The two hybrid-origin candidates required distinct inferential strategies. Sampled-parent hybrid origin in *A. gambiae* was evaluated through temporal equivalence between reticulation and lineage origin, whereas the ghost-mediated origin of *A. merus* was resolved by direct marginal-likelihood comparison of alternative network histories. Together, these analyses extend hybrid-origin inference to both sampled-parent and ghost-mediated histories.

### Parental-affinity loci suggest a route to reproductive isolation in *A. gambiae*

The 3L and 3R candidate regions identify parental genomic components that may have contributed to the subsequent establishment or maintenance of reproductive isolation in *A. gambiae*. The *A. coluzzii*-affinity 3L interval contains OBP4, Or43, and Or44, placing parental ancestry within an olfactory gene set involved in odor detection (Carey et al. 2010; Wang et al. 2010; Davrazou et al. 2011; Qiao et al. 2011). Gene trees independently recover the same *A. gambiae*-*A. coluzzii* affinity at these loci.

The *A. arabiensis*-affinity 3R interval contains the RR-2 cuticular-protein cluster CPR82-CPR100. RR-2 proteins are structural components of the insect cuticle, and AGAP010105 independently groups *A. gambiae* with *A. arabiensis* (Cornman et al. 2008; Togawa et al. 2008; Vannini and Willis 2017). The contrasting affinities of the 3L olfactory region and the 3R cuticular region therefore identify a mosaic of parental genomic components in biological systems with potential roles in prezygotic isolation.

This genomic pattern parallels the mechanism demonstrated in *Ostryopsis intermedia*, where a hybrid lineage inherited alternate isolating alleles from its parental species (Wang et al. 2021). In *A. gambiae*, our results generate a comparable but testable mechanistic hypothesis: hybridization brought together parental variants in olfactory and cuticular systems that may subsequently have contributed to reproductive isolation and the maintenance of lineage independence. Ligand-response and behavioral assays of the 3L genes, expression and structural analyses of the 3R cluster, and ancestry-informed crosses combining the two regions can directly test this hypothesis.

## Materials and Methods

### Genome resources, gene-set curation, and orthogroup inference

We obtained independently annotated genome assemblies and gene sets for six species in the *A. gambiae* complex: *A. gambiae* (Agam), *A. coluzzii* (Acol), *A. arabiensis* (Aara), *A. melas* (Amel), *A. merus* (Amer), and *A. quadriannulatus* (Aqua). *A. christyi* and *A. epiroticus* were included as outgroups. All genome assemblies and associated protein-coding annotations were retrieved from public biological databases and download links are provided in table S19.

To reduce annotation-derived heterogeneity among species, all gene sets were processed with a common annotation-curation pipeline. GFF3 files were standardized and repaired with AGAT (Dainat 2024). For genes with multiple transcript isoforms, only the longest protein-coding isoform was retained. Redundant overlapping gene models were then removed at the transcript level. Coding-sequence coordinates were used to define the genomic span of each retained transcript, and BEDTools (Quinlan and Hall 2010) was used to identify overlapping spans on the same scaffold or chromosome. Overlapping transcripts were grouped into connected components, and a single representative was retained from each component, prioritizing the transcript encoding the longest protein sequence. Coding and protein sequences were extracted with gffread (Pertea and Pertea 2020). Transcripts were retained only when the coding-sequence length was divisible by three and the translated protein contained no internal stop codons.

To evaluate sensitivity to short or potentially fragmented gene predictions, SeqKit (Shen et al. 2016) was used to generate four protein datasets with different minimum-length thresholds: no additional cutoff, ≥50 amino acids, ≥100 amino acids, and ≥150 amino acids. Orthogroups were inferred independently for each dataset with OrthoFinder (Emms and Kelly 2019) using its default DIAMOND-based sequence-search procedure.

### Gene-content species-tree inference and robustness analyses

For each orthogroup dataset, we constructed a binary presence-absence matrix with species as rows and orthogroups as columns. An orthogroup was coded as present when at least one gene copy was assigned to that species and absent otherwise. Multi-copy orthogroups were collapsed to a single presence state to prevent lineage-specific duplications from receiving disproportionate weight. Orthogroups present in all taxa were removed because they were uninformative for presence-absence phylogenetic inference.

Species trees were inferred from the binary matrices with IQ-TREE 2 (Minh et al. 2020) under the Mk+R+FO model with ascertainment-bias correction (+ASC). Nodal support was assessed with 1,000 nonparametric bootstrap replicates. Trees were rooted with *A. epiroticus*. The resulting gene-content trees were treated as candidate fixed species-tree hypotheses in downstream D-BPP analyses.

Robustness to stochastic error in presence-absence coding was evaluated by perturbing the primary ≥100-amino-acid matrix. Each binary entry xij∈{0,1} was independently flipped with probability p, such that xij′=1−xij with probability p and xij′=xij otherwise. This procedure introduced symmetric false-gain and false-loss errors. We evaluated p=0.001, 0.005, and 0.01 and generated 10 independent replicates at each perturbation level using different random seeds. Trees were re-inferred from every perturbed matrix using the same IQ-TREE 2 settings as for the unperturbed data.

### Noncoding loci and sequence-based species-tree inference

For *D*-statistic screening and BPP analyses, we used the publicly available multilocus noncoding sequence dataset compiled by Thawornwattana et al. (2018) from the whole-genome alignments originally generated by Fontaine et al. (2015). The realigned loci and accompanying processing scripts provided by Thawornwattana et al. (2018) were downloaded and subjected to additional filtering for the present analyses. We retained alignments 100 to 1,000 base pairs long, separated adjacent loci by at least 20 kb to reduce linkage, and excluded alignments containing more than 50% missing or gap characters across the included taxa. Because previous studies have shown that sex-linked regions and known inversion regions in the *A. gambiae* complex can follow evolutionary histories distinct from those of the broader autosomal genome (Fontaine et al. 2015; Thawornwattana et al. 2018), loci in putatively sex-linked regions or overlapping known inversion regions were excluded to reduce the influence of region-specific histories on downstream analyses. The final dataset contained 4,214 noncoding loci and was used for all *D*-statistic and BPP analyses.

For sequence-based species-tree inference, a locus tree was estimated independently for each alignment with IQ-TREE 2. The best-fitting nucleotide-substitution model was selected using the program’s automatic model-selection procedure, and branch support was assessed with 1,000 ultrafast bootstrap replicates. Because *A. epiroticus* was absent from this dataset, locus trees were rooted with *A. christyi*. The rooted locus trees were summarized with ASTRAL-III (Zhang et al. 2018) to obtain a coalescent-based sequence species tree, which was evaluated as an alternative species-tree hypothesis in the D-BPP analyses (fig. S2).

### Simulation-based assessment of model misspecification

We evaluated how misspecification of reticulation-event order, placement, or mode affected posterior time estimates under four reticulate-history scenarios (Fig. 3): Model 1, two introgression events from a sampled donor; Model 2, two introgression events from a ghost lineage; Model 3, hybrid speciation with both parental lineages sampled; and Model 4, ghost-mediated hybrid speciation. Multilocus alignments were generated with the BPP simulation module (Yang 2015; Flouri et al. 2018) under JC69, with two sequences per sampled species, 1,000 loci, and 1,000 bp per locus.

The population-size parameter was fixed at θ=0.01, and divergence and reticulation times were parameterized relative to θ. In BPP, τ and θ are measured in expected substitutions per site, and one coalescent unit corresponds to θ/2.

For Model 1, parameters were τR=2θ, τX=θ, τM=τN=0.75θ, and τP=τQ=0.5θ, with introgression probability φ1=0.2 and φ2=0.4. For Model 2, parameters were τS=3θ, τR =2θ, τX=θ, τM=τN=0.75θ, and τP=τQ=0.5θ, with φ1=0.2 and φ2=0.4. For Model 3, parameters were τR=2θ and τX=τN=τM=θ, with parental contributions φ1=0.6 and φ2 =0.4. For Model 4, all sequences from parental lineage C were removed from the Model 3 datasets, thereby treating C as an unsampled ghost parent.

Each simulated dataset was analyzed under the generating model and one or more misspecified alternatives. For Model 1, the misspecified model reversed the order of the two introgression events. For Model 2, we evaluated both an incorrect event order and assignment of ghost introgression to an incorrect ancestral branch. For Model 3, data generated under hybrid speciation were analyzed under an ordinary post-divergence introgression model. For Model 4, ghost-mediated hybrid-speciation data were analyzed under a post-divergence ghost-introgression model. Ten independent replicates were analyzed for each comparison. Posterior means and 95% highest posterior density intervals were compared with the generating values to quantify bias in inferred reticulation timing, event placement, and evolutionary interpretation.

For these BPP analyses, the population-size parameter was assigned a gamma prior, θ∼G(2,200), with prior mean 0.01, and the root-age parameter was assigned G(2,67), with prior mean approximately 0.03. Here, G(α,β) denotes a gamma distribution parameterized by shape α and rate β. Each analysis used a burn-in of 6 × 10^4^ iterations, followed by 2 × 10^5^ posterior samples recorded every two iterations.

### Simulations evaluate the Savage-Dickey density-ratio test for hybrid origin

Candidate hybrid-speciation events were evaluated by comparing the posterior distributions of the relevant species-divergence time (τX) and reticulation time (τN). Post-divergence introgression requires τN < τX, whereas a hybrid-origin scenario predicts temporal equivalence between hybridization and lineage formation. We therefore tested H0:τX=τN against H1:τN<τX using the Savage-Dickey density-ratio procedure applied by Thawornwattana et al. (2025).

Because exact equality has zero probability under continuous prior and posterior distributions, temporal equivalence was represented by the interval 0<τX−τN<ε. We evaluated ε=10^−5^ and 10^−6^ and calculated Bε=P(Eε)/ P(Eε∣X), where Eε denotes the temporal-equivalence interval, P(Eε) is its prior probability, and P(Eε∣X) is its posterior probability. Prior probabilities were estimated from additional BPP MCMC analyses conducted without sequence data (usedata = 0), whereas posterior probabilities were obtained from the corresponding analyses using sequence data (usedata = 1). All model parameters and prior settings were kept identical between the prior and posterior analyses. For each value of ε, the two probabilities were estimated as the proportions of MCMC samples satisfying 0<τX−τN<ε. Following Thawornwattana et al. (2025), Bε<0.01 was interpreted as support for the equal-time hybrid-speciation model, whereas Bε>100 supported the post-divergence introgression model. Values between these thresholds were treated as inconclusive.

The performance of this test was evaluated using datasets of 5,000 loci simulated under Models 3 and 4, with all other simulation settings as described above. Ten independent replicate datasets were generated for each scenario. We applied the Savage-Dickey test under two scenarios: one in which both parental lineages were sampled and another in which one parental lineage was a ghost lineage (extinct or unsampled), to evaluate the behavior of the temporal-equivalence test under sampled-parent and ghost-mediated hybrid-speciation histories.

### D-BPP network inference under alternative species-tree hypotheses

For each alternative species tree, we used the D-BPP framework (Yang et al. 2026), which couples the speed of genome-wide *D*-statistics with the accuracy of Bayesian full-likelihood inference to infer networks involving both extant and ghost introgression. Within the D-BPP workflow, *D*-statistic calculations were performed as follows: concatenated sequence alignments were first converted to VCF using SNP-sites (Page et al. 2016), and *D*-statistics were subsequently computed with Dsuite (Malinsky et al. 2021). For each significant triple (*p* < 0.01, after Bonferroni correction for multiple testing), we calculated 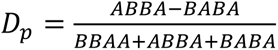 (Hamlin et al. 2020) and ranked the resulting values in decreasing order to prioritize candidate introgression events. We then removed outgroup sequences for subsequent BPP analysis. Each significant triple was evaluated under three candidate histories: introgression from P3 to P2, introgression from P2 to P3, and ghost introgression to P1. The two directional events between sampled lineages were evaluated within a bidirectional-introgression model following the D-BPP framework (Yang et al. 2026), whereas ghost introgression was evaluated separately. Candidate introgression events were sequentially evaluated using the Bayesian test implemented in BPP (Flouri et al. 2020; Ji et al. 2023). The test compares a model containing the candidate introgression event (H1) with the corresponding no-introgression model (H0) using a Savage-Dickey density-ratio approximation. Because the no-introgression condition corresponds to an inheritance probability φ=0, which lies on the boundary of the parameter space, a small null interval φ<ε within the H1 model was used to approximate H0. The Bayes factor was approximated as B10,ε= P(ϕ<ε)/P(ϕ<ε∣X), with the numerator and denominator representing the prior and posterior probabilities of the null interval, respectively. As ε→0, B10,ε converges to the Bayes factor B10.

Following Ji et al. (2023), we used ε=0.01 and confirmed that ε=0.001 yielded similar results. Candidate events with B10>100 were considered decisively supported and were retained during network construction.

When multiple candidate events explained overlapping triples, we applied a parsimony criterion that favored the smallest set of reticulation edges accounting for the largest number of significant signals. Networks were constructed iteratively, beginning with the highest-ranked *D*p signals. Reticulation events receiving decisive Bayesian support were incorporated into the network, triples accounted for by the retained events were removed, and the procedure was repeated for the remaining unexplained signals. This process continued until the significant signals associated with each starting topology had either been incorporated into the network or explicitly evaluated and rejected. Detailed reconstruction steps for the alternative topologies are provided in Supplementary Texts S1 to S5.

### Marginal-likelihood comparison of topology-network models

Marginal likelihoods were estimated in BPP by thermodynamic integration with Gaussian quadrature using 16 integration points (Lartillot and Philippe 2006; Rannala and Yang 2017). For model comparison, the 4,214 loci were divided into four nonoverlapping partitions containing 1,057, 1,050, 1,057, and 1,050 loci. Loci were first separated into odd- and even-indexed sets according to their order in the curated dataset, and each set was then divided sequentially into two subsets. The same four data partitions were used for all marginal-likelihood comparisons described below.

Under the gene-content species-tree topology, we first compared two alternative reticulate histories. In the first model, the ghost contributions associated with *A. merus* and *A. melas* were represented as separate post-divergence introgression events. In the alternative model, *A. merus* was modeled as a ghost-mediated hybrid lineage, whereas the ghost contribution to *A. melas* remained a post-divergence introgression event. All other retained reticulation events were identical between the two models. The two models were evaluated independently for each of the four data partitions, and the better-supported model was retained as the completed network associated with the gene-content species-tree topology.

We then compared the completed D-BPP networks constructed on the five alternative species-tree hypotheses, including the retained gene-content topology-network model and the networks reconstructed under the four sequence-based species-tree hypotheses. Each topology-network model was evaluated independently using the same four data partitions, and model rankings were compared across partitions to assess the consistency of support among alternative reticulate histories.

For marginal-likelihood estimation, the population-size parameter was assigned a prior of θ∼IG(3,0.002), and the root-age parameter was assigned τ0∼IG(3,0.01), following the parameterization implemented in BPP. At each of the 16 integration points, the MCMC was run with a burn-in of 1 × 10^6^ iterations, followed by 1 × 10^7^ iterations, with samples recorded every two iterations. All competing models were analyzed using identical prior distributions and MCMC settings within each data partition.

After identifying the best-supported topology-network model, its posterior parameters were estimated using all 4,214 loci. The population-size parameter was assigned θ∼G(2,200), with prior mean 0.01, and the root-age parameter was assigned τ0 ∼G(2,67), with prior mean approximately 0.03. The MCMC used a burn-in of 1 × 10^5^ iterations, followed by 1.5 × 10^6^ iterations, with samples recorded every two iterations. Posterior estimates of divergence and reticulation times were converted to absolute time using a mutation rate of 2.8 × 10^−9^ substitutions per site per generation and 11 generations per year, using the mutation rate adopted by Thawornwattana et al. (2018), based on Keightley et al. (2014). Absolute ages are reported in millions of years ago (Ma).

### Simulation under the empirically best-supported network

We additionally simulated multilocus data under the best-supported empirical MSci network to assess whether D-BPP could recover a reticulate history of comparable complexity. The simulation included six ingroup lineages corresponding to *A. melas*, *A. merus*, *A. gambiae*, *A. coluzzii*, *A. arabiensis*, and *A. quadriannulatus*, together with one outgroup. At each locus, two sequences were sampled from each ingroup species and one sequence from the outgroup. Posterior mean estimates of divergence and reticulation times (τ), population-size parameters (θ), and introgression probability (φ) obtained under the best-supported empirical network were used as the generating parameter values for the ingroup network. For the outgroup, the divergence-time and population-size parameters were fixed at τ=0.08 and θ=0.179, respectively, based on the estimates reported in Fig. 2A of Thawornwattana et al. (2018). Using JC69 in BPP, we generated 4,214 independent loci of 311 bp, matching the number and mean length of the empirical noncoding loci. *D*-statistics were first calculated with the outgroup included, after which the outgroup was removed and Bayesian event testing and network construction were performed in BPP following the same D-BPP workflow used for the empirical data. Recovery was assessed by comparing the inferred reticulation edges with those in the generating network.

### Temporal tests of the hybrid origin of *A. gambiae*

Within the best-supported full network, we first tested whether the retained *A. arabiensis* → *A. gambiae* reticulation coincided with the divergence of *A. gambiae* and *A. coluzzii*. The Savage-Dickey temporal-equivalence test described above was applied to the corresponding reticulation time (τN) and lineage-divergence time (τX), with temporal equivalence represented by 0 < τX − τN < ε for ε = 10^−5^ and 10^−6^. Prior probabilities of the temporal-equivalence interval were estimated from matched BPP MCMC runs without sequence data (usedata = 0), using the same full-network structure and prior settings as the posterior analyses. Posterior probabilities were estimated from the corresponding analyses with sequence data (usedata = 1), and Bε was calculated as the ratio of the prior to posterior probability of the interval.

We then evaluated the same hybrid-origin event in a reduced model containing *A. coluzzii*, *A. gambiae*, and *A. arabiensis*. The local branching relationship among these species was identical across the five candidate species-tree hypotheses, allowing the temporal relationship between reticulation and lineage divergence to be evaluated without reticulation events elsewhere in the network. The reduced model retained the three reticulation events supported among these lineages: introgression from *A. arabiensis* to *A. gambiae*, introgression from *A. gambiae* into *A. arabiensis*, and ghost introgression into *A. coluzzii*. The two directional events between *A. gambiae* and *A. arabiensis* were modeled independently with separate, unconstrained time parameters. The same Savage-Dickey procedure was used to test temporal equivalence between the *A. arabiensis* → *A. gambiae* reticulation time and the *A. gambiae-A. coluzzii* divergence time.

To assess the stability of this temporal-equivalence signal under the empirical parameter regime and data scale, we simulated 10 replicate datasets using posterior mean parameter estimates from the reduced three-species MSci network. Each replicate contained three ingroup lineages corresponding to *A. gambiae*, *A. coluzzii*, and *A. arabiensis*, with two sequences sampled per species at each locus, and 4,214 independent loci of 311 bp, matching the number and mean length of the empirical noncoding loci. Sequences were simulated under JC69 in BPP. The same Savage-Dickey test was applied to each replicate to assess temporal equivalence between the *A. arabiensis* → *A. gambiae* reticulation time and the *A. gambiae-A. coluzzii* divergence time.

### Population genomic variant data

Population genomic variants in VCF format were obtained from the Ag3.0 release of the *Anopheles gambiae* 1000 Genomes Project (Ag1000G phase 3) (The *Anopheles gambiae* 1000 Genomes Consortium 2021). We used individuals from the AG1000G-BF-B, AG1000G-CM-C, AG1000G-KE, and AG1000G-MW cohorts, representing *A. gambiae*, *A. coluzzii*, and *A. arabiensis*. Species assignments followed the metadata provided with the Ag1000G phase 3 release. To balance sample sizes among species, 59 individuals were retained for each species, yielding 177 individuals in total.

Sample IDs and species assignments for all retained individuals are provided in table S20. The downloaded Ag1000G phase 3 VCF files were merged with bcftools and subset to the selected individuals but were otherwise used without additional variant-level filtering. These balanced population datasets were used for the genome-wide selection analyses and for reconstruction of individual consensus sequences used in the candidate-gene phylogenetic analyses.

### Genome-wide XP-CLR scans

Cross-Population Composite Likelihood Ratio (XP-CLR) (Chen et al. 2010) scans were conducted with *A. gambiae* as the target population and each putative parental species as a separate reference population. The two comparisons were *A. gambiae* versus *A. arabiensis* and *A. gambiae* versus *A. coluzzii*. Genetic distances were approximated from the recombination map used in the analysis, and XP-CLR scores were summarized in 20-kb windows. Windows falling within the upper 0.5% of the genome-wide XP-CLR score distribution for each comparison were designated as candidate windows. Candidate windows separated by less than 1 Mb were merged with BEDTools (Quinlan and Hall 2010) to define continuous candidate intervals.

### Comparison-specific intervals and parental-ancestry assignment

To identify genomic regions showing comparison-specific differentiation, candidate intervals overlapping between the two XP-CLR scans were excluded. Intervals detected only in the *A. gambiae*-versus-*A. coluzzii* comparison were classified as showing greater affinity of *A. gambiae* to *A. arabiensis*. Conversely, intervals detected only in the *A. gambiae*-versus-*A. arabiensis* comparison were classified as showing greater affinity of *A. gambiae* to *A. coluzzii*. These comparison-specific patterns were used as regional parental-affinity assignments for subsequent HKA validation.

### HKA validation of comparison-specific XP-CLR intervals

Comparison-specific intervals were further evaluated using HKA tests (Hudson et al. 1987). For each interval, we counted the numbers of segregating sites (S) and fixed differences (D) from the joint VCF file. S was quantified as the number of polymorphic sites segregating within the focal population, with loci containing more than 20% missing data excluded. D was defined as the number of sites at which the two compared species were reciprocally fixed for alternative alleles. We carried out the HKA test by comparing the S/D ratio for each candidate interval with the background ratio, which was computed with the sum of S and D values across all analyzed intervals. A Fisher’s exact test on the 2×2 contingency table was used to test the null hypothesis that S(candidate interval)/D(candidate interval)=S(genomic background)/D(genomic background). A candidate interval with reduced S/D ratio was considered positively selected and a Bonferroni-corrected P < 0.05 was considered significant. An interval was retained only when the HKA criterion was satisfied in the putative source lineage inferred from the comparison-specific XP-CLR pattern.

### Gene annotation and locus-specific ancestry assessment

Genes overlapping the retained XP-CLR and HKA-supported intervals were extracted from the AgamP4 reference annotation and examined for functions potentially relevant to prezygotic reproductive isolation. Functional interpretation was based on the reference gene annotation and published experimental evidence. For informative candidate genes, individual consensus sequences were reconstructed from the joint VCF using *bcftools consensus*, guided by the coordinates of the AgamP4 reference genome. The resulting sequences from *A. gambiae*, *A. coluzzii*, and *A. arabiensis* were used for locus-specific phylogenetic reconstruction. Gene trees were reconstructed in IQ-TREE 3 (Wong et al. 2026) using the BIONJ method under the Kimura two-parameter (K2P) nucleotide model. The resulting gene trees were used as an independent locus-level assessment of whether *A. gambiae* showed greater sequence affinity to *A. arabiensis* or *A. coluzzii* within the candidate regions.

## Acknowledgments

We acknowledge the genomic resources and whole-genome alignments of the *Anopheles gambiae* species complex generated by Fontaine et al. (2015), and the multilocus sequence datasets subsequently compiled and processed by Thawornwattana et al. (2018), which formed the basis of our *D*-statistic screening and BPP analyses. We also thank the *Anopheles gambiae* 1000 Genomes (Ag1000G) Consortium and its contributing investigators for generating and making available through MalariaGEN the Ag3.0 (Ag1000G phase 3) genomic data used in the population genomic analyses. We acknowledge the use of ChatGPT (OpenAI) to assist with English-language editing and stylistic refinement. All scientific content, analyses, interpretations, and conclusions were reviewed and verified by the authors.

## Author contributions

DYZ and WNB conceived the study. BWZ and YY curated the data. YY and XXP performed the formal analyses. BWZ and DYZ acquired funding. YY, XXP, and WNB developed the methodology. WNB, BWZ, and DYZ supervised the study. YY prepared the visualizations. YY, BWZ, and DYZ wrote the original draft, and XXP and WNB reviewed and edited the manuscript.

## Data availability

All genome assemblies, genome annotations, sequence alignments, and population genomic data analyzed in this study were obtained from publicly available databases or previously published datasets. Genome assembly and annotation resources are listed in table S19. Population genomic variant data were obtained from the MalariaGEN Ag3.0 (Ag1000G phase 3) resource. The population genomic selection analyses included 177 individuals, comprising 59 individuals each of *A. gambiae*, *A. coluzzii*, and *A. arabiensis*. The D-BPP workflow and associated analysis scripts are publicly available at https://github.com/yangyang9608/D-BPP_Workflow.

