## Supplementary material for "Hybrid speciation and ghost ancestry shape the *Anopheles gambiae* species complex": fig. S1; table S1

### Supplementary Information

▪ **Supplementary Texts**

**Table S20. Sample IDs of the Ag1000G phase 3 individuals used in population genomic analyses. Each species is represented by 59 individuals36**

### Text S1. D-BPP analysis under the gene-family content species tree

For the gene-family content species tree with *A. christyi* as the outgroup, *D*-statistics identified 13 significant taxon trios after multiple-testing correction (adjusted $p < 0.01$; table S3). Following the principle of parsimony, we focused on the six highest-ranking significant trios based on *D*p values, all of which involved the *A. merus*–*A. melas* sister clade (P1), the three-species clade comprising *A. gambiae*, *A. coluzzii*, and *A. arabiensis* (P2), and *A. quadriannulatus* (P3). Accordingly, we tested three candidate introgression events that could parsimoniously explain these trios: inflow from P3 to P2 (P3 → P2), outflow from P2 to P3 (P2 → P3), and introgression from an unsampled ghost lineage into P1 (Ghost → P1). Bayesian testing provided decisive support for the inflow and ghost-introgression models ($B_{10} > 100$). However, the estimated time of ghost introgression to the ancestral lineage of *A. merus* and *A. melas* was very close to their divergence time. Our simulations showed that such temporal coincidence can arise artifactually when ghost introgression is assigned to an incorrect ancestral branch (Fig. 3). We therefore tested an alternative hypothesis involving two lineage-specific ghost-introgression events, one to *A. merus* and the other to *A. melas*. Both events were strongly supported by Bayesian testing in BPP ($B_{10}$>100). This two-event hypothesis also accounted for the next four significant D-statistic signals involving the *A. merus*–*A. melas* pair (table S3; Amer–Amel–Acol, Amer–Amel–Agam, Amer–Amel–Aara, and Amer–Amel–Aqua). We next evaluated the 11$^{th}$ significant trio, Acol-Agam-Aara, involving the three principal vector species. One signal was compatible with three alternative reticulation events: inflow and outflow between *A. gambiae* and *A. arabiensis*, and ghost introgression to *A. coluzzii*. Bayesian tests supported all three candidate models ($B_{10} > 100$; Fig. S3; Table S4). For the final two significant trios (Table S3; Aara-Agam-Aqua and Aara-Acol-Aqua), we tested two alternative gene flow events: inflow from *A. quadriannulatus* into the ancestral lineage of *A. gambiae* and *A. coluzzii*, or outflow with reverse direction. Bayesian testing in BPP supported only the outflow event.

Under the gene-family content species-tree, after all retained reticulation events had been incorporated into the completed network, the posterior estimates continued to place the ghost-lineage contribution to A. merus very close to the divergence of A. merus and A. melas (table S9). This temporal proximity suggested a possible ghost-mediated hybrid origin of A. merus. However, because our simulations showed that timing patterns alone are insufficient to distinguish ghost-mediated hybrid speciation from post-divergence ghost introgression, we compared two completed MSci network models using marginal likelihoods. In the first model, the ghost-lineage contributions to both A. merus and A. melas were modeled as post-divergence introgression. In the alternative model, A. merus was modeled as a ghost-mediated hybrid species, whereas A. melas retained a post-divergence ghost-introgression event. All other speciation and reticulation events were held constant between the two models. The network incorporating a ghost-mediated hybrid origin of A. merus had the higher marginal likelihood, supporting this interpretation over the alternative post-divergence ghost-introgression model (table S10).

### Text S2. D-BPP analysis under the sequence-based species tree

We applied the D-BPP framework (Yang et al. 2026) to the sequence-based species tree and identified 10 significant *D*-statistic triples (table S5). Following the parsimony criterion described above, the first four significant triples were jointly evaluated by testing introgression from the ancestral lineage of the four-species clade comprising *A. quadriannulatus*, *A. gambiae*, *A. coluzzii*, and *A. arabiensis* to *A. merus*, introgression in the reverse direction, and ghost introgression to *A. melas*. Bayesian testing decisively supported ghost introgression to *A. melas* and introgression from the four-species ancestral lineage to *A. merus* ($B_{10} > 100$; table S4; fig. S3C).

The fifth significant triple involved *A. gambiae*, *A. arabiensis*, and *A. coluzzii*. We tested introgression from *A. gambiae* to *A. arabiensis*, introgression from *A. arabiensis* to *A. gambiae*, and ghost introgression to *A. coluzzii*. All three events received decisive support ($B_{10} > 100$; table S4; fig. S3C) and were retained in the network.

The sixth and seventh significant triples were then jointly evaluated by testing introgression from *A. quadriannulatus* to the ancestral lineage of *A. gambiae* and *A. coluzzii*, introgression in the reverse direction, and ghost introgression to *A. arabiensis*. Bayesian testing decisively supported introgression from the *A. gambiae*–*A. coluzzii* ancestral lineage to *A. quadriannulatus* and ghost introgression to *A. arabiensis* ($B_{10} > 100$; table S4; fig. S3C).

Finally, after incorporating the supported reticulation events described above, the eighth to tenth significant triples remained unexplained and were evaluated by testing bidirectional gene flow between *A. quadriannulatus* and *A. merus*. Neither direction was supported by Bayesian testing in BPP ($B_{10} = 0.01$ for both; table S4; fig. S3C), and neither event was retained. Accordingly, these three significant D-statistic triples remained unexplained in the final network.

Thus, the final network under the sequence-based species tree contained seven reticulation events: (i) ghost introgression to *A. melas*; (ii) introgression from the ancestral lineage of *A. quadriannulatus*, *A. gambiae*, *A. coluzzii*, and *A. arabiensis* to *A. merus*; (iii) introgression from *A. gambiae* to *A. arabiensis*; (iv) introgression from *A. arabiensis* to *A. gambiae*; (v) ghost introgression to *A. coluzzii*; (vi) introgression from the ancestral lineage of *A. gambiae* and *A. coluzzii* to *A. quadriannulatus*; and (vii) ghost introgression to *A. arabiensis* (fig. S3C).

### Text S3. D-BPP analysis under the species topology of Thawornwattana et al. (2018)

We applied D-BPP to the species topology reported by Thawornwattana et al. (2018) and identified 14 significant *D*-statistic triples (table S6). Following the parsimony criterion described above, the first 10 significant triples were jointly evaluated by testing introgression from the ancestral lineage of *A. gambiae* and *A. coluzzii* to the ancestral lineage of *A. arabiensis* and *A. quadriannulatus*, introgression in the reverse direction, and ghost introgression to *A. melas*. Bayesian testing decisively supported ghost introgression to *A. melas* and introgression from the *A. gambiae*–*A. coluzzii* ancestral lineage to the *A. arabiensis*–*A. quadriannulatus* ancestral lineage ($B_{10} > 100$; table S4; fig. S3D). These two retained events jointly accounted for the first 10 significant triples.

The 11th significant triple involved *A. gambiae*, *A. arabiensis*, and *A. coluzzii*. We tested introgression from *A. gambiae* to *A. arabiensis* and introgression from *A. arabiensis* to *A. gambiae*. Ghost introgression to *A. coluzzii* was not evaluated because, under this topology, such an event would also be expected to generate a significant *A. coluzzii*–*A. gambiae*–*A. quadriannulatus* triple, which was not observed. Both directions of introgression between *A. gambiae* and *A. arabiensis* received decisive Bayesian support ($B_{10} > 100$; table S4; fig. S3D) and were retained.

The 12th to 14th significant triples all involved *A. quadriannulatus* and *A. merus* and were therefore jointly evaluated by testing bidirectional introgression between these two species. Introgression from *A. quadriannulatus* to *A. merus* received decisive Bayesian support, whereas the reverse direction was not supported (table S4; fig. S3D). The supported event was retained in the final network. Thus, all 14 significant *D*-statistic triples were accounted for by the retained reticulation events.

The final network under the Thawornwattana et al. (2018) topology contained five reticulation events: (i) ghost introgression to *A. melas*; (ii) introgression from the ancestral lineage of *A. gambiae* and *A. coluzzii* to the ancestral lineage of *A. arabiensis* and *A. quadriannulatus*; (iii) introgression from *A. arabiensis* to *A. gambiae*; (iv) introgression from *A. gambiae* to *A. arabiensis*; and (v) introgression from *A. quadriannulatus* to *A. merus* (fig. S3D).

### Text S4. D-BPP analysis under the X-chromosome topology of Fontaine et al. (2015)

We applied D-BPP to the X-chromosome topology reported by Fontaine et al. (2015) and identified 16 significant *D*-statistic triples (table S7). Following the parsimony criterion described above, the first two significant triples were jointly evaluated by testing introgression from *A. arabiensis* to the ancestral lineage of *A. gambiae* and *A. coluzzii*, introgression in the reverse direction, and ghost introgression to *A. merus*. Bayesian testing decisively supported ghost introgression to *A. merus* and introgression from *A. arabiensis* to the *A. gambiae*–*A. coluzzii* ancestral lineage ($B_{10}$ > 100; table S4; fig. S3E). Incorporation of these two events explained the next eight significant triples (the third to tenth), which were therefore removed from further testing.

The remaining significant triples were evaluated according to their corresponding parsimonious gene-flow scenarios. The 11th triple was evaluated by testing bidirectional introgression between *A. quadriannulatus* and *A. merus*. The 12th and 13th triples were jointly evaluated by testing bidirectional introgression between *A. melas* and the ancestral lineage of *A. gambiae* and *A. coluzzii*. The 14th triple was evaluated by testing bidirectional introgression between *A. arabiensis* and *A. merus*, and the 15th by testing bidirectional introgression between *A. gambiae* and *A. arabiensis*. The 16th triple corresponded to the same *A. quadriannulatus*–*A. merus* gene-flow scenario already evaluated for the 11th triple and therefore required no additional test. Additional ghost-introgression models were not evaluated for these residual signals because, under the corresponding topology, such ghost events would be expected to generate additional associated significant triples that were not observed.

Among these candidate events, only the two directional introgression events between *A. gambiae* and *A. arabiensis* received decisive Bayesian support ($B_{10}$ > 100; table S4; fig. S3E), and both were retained. None of the other candidate gene-flow events received decisive support and they were therefore excluded from the final network. Consequently, five of the 16 significant *D*-statistic triples remained unexplained by the final network.

The final network under the Fontaine et al. (2015) X-chromosome topology contained four reticulation events: (i) ghost introgression to *A. merus*; (ii) introgression from *A. arabiensis* to the ancestral lineage of *A. gambiae* and *A. coluzzii*; (iii) introgression from *A. gambiae* to *A. arabiensis*; and (iv) introgression from *A. arabiensis* to *A. gambiae* (fig. S3E).

### Text S5. D-BPP analysis under the whole-genome topology of Fontaine et al. (2015)

We applied D-BPP to the whole-genome topology reported by Fontaine et al. (2015) and identified 10 significant *D*-statistic triples (table S8). Following the parsimony criterion described above, the first four significant triples were jointly evaluated by testing introgression from the ancestral lineage of the four-species clade comprising *A. quadriannulatus*, *A. gambiae*, *A. coluzzii*, and *A. arabiensis* to *A. melas*, introgression in the reverse direction, and ghost introgression to *A. merus*. Bayesian testing decisively supported ghost introgression to *A. merus* and introgression from the four-species ancestral lineage to *A. melas* ($B_{10}$ > 100; table S4; fig. S3F).

The fifth significant triple involved *A. gambiae*, *A. arabiensis*, and *A. coluzzii*. We tested introgression from *A. gambiae* to *A. arabiensis*, introgression from *A. arabiensis* to *A. gambiae*, and ghost introgression to *A. coluzzii*. All three events received decisive support ($B_{10}$ > 100; table S4; fig. S3F) and were retained in the network.

The sixth and seventh significant triples were then jointly evaluated by testing introgression from *A. quadriannulatus* to the ancestral lineage of *A. gambiae* and *A. coluzzii*, introgression in the reverse direction, and ghost introgression to *A. arabiensis*. Bayesian testing decisively supported introgression from the *A. gambiae*–*A. coluzzii* ancestral lineage to *A. quadriannulatus* and ghost introgression to *A. arabiensis* ($B_{10}$ > 100; table S4; fig. S3F), whereas introgression in the reverse direction was not supported.

Finally, after incorporating the supported reticulation events described above, the eighth to tenth significant triples remained and all involved *A. quadriannulatus* and *A. merus*. Following the parsimony criterion, these three triples were jointly evaluated by testing bidirectional introgression between the two species. Introgression from *A. quadriannulatus* to *A. merus* received decisive Bayesian support ($B_{10}$ > 100), whereas the reverse direction was not supported ($B_{10}$ = 0.01; table S4; fig. S3F). The supported event was therefore retained in the final network. Thus, all 10 significant *D*-statistic triples were accounted for by the retained reticulation events.

The final network under the Fontaine et al. (2015) whole-genome topology contained eight reticulation events: (i) ghost introgression to *A. merus*; (ii) introgression from the ancestral lineage of *A. quadriannulatus*, *A. gambiae*, *A. coluzzii*, and *A. arabiensis* to *A. melas*; (iii) introgression from *A. gambiae* to *A. arabiensis*; (iv) introgression from *A. arabiensis* to *A. gambiae*; (v) ghost introgression to *A. coluzzii*; (vi) introgression from the ancestral lineage of *A. gambiae* and *A. coluzzii* to *A. quadriannulatus*; (vii) ghost introgression to *A. arabiensis*; and (viii) introgression from *A. quadriannulatus* to *A. merus* (fig. S3F).

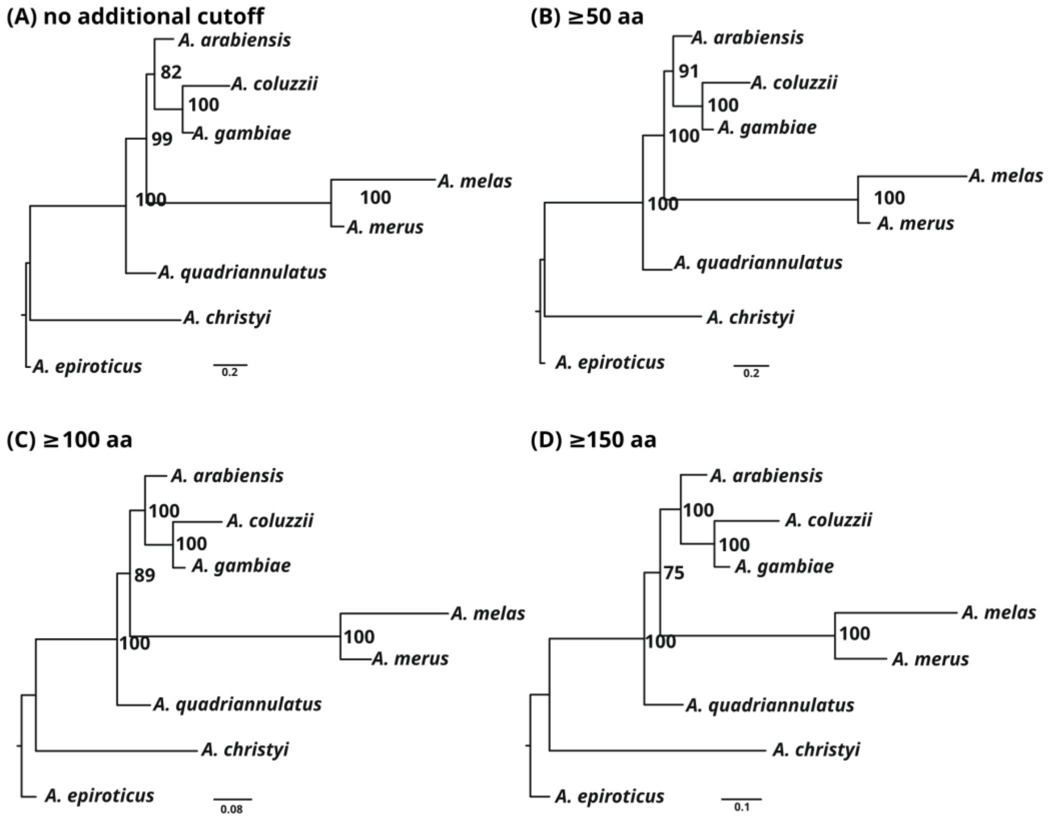

**Fig. S1.**
Gene-family content phylogenies inferred under different minimum protein-length thresholds. Gene-family content phylogenies were inferred from orthogroup presence/absence matrices constructed using four filtering schemes: (A) no additional protein-length cutoff, (B) ≥50 aa, (C) ≥100 aa, and (D) ≥150 aa. Node labels indicate bootstrap support values.

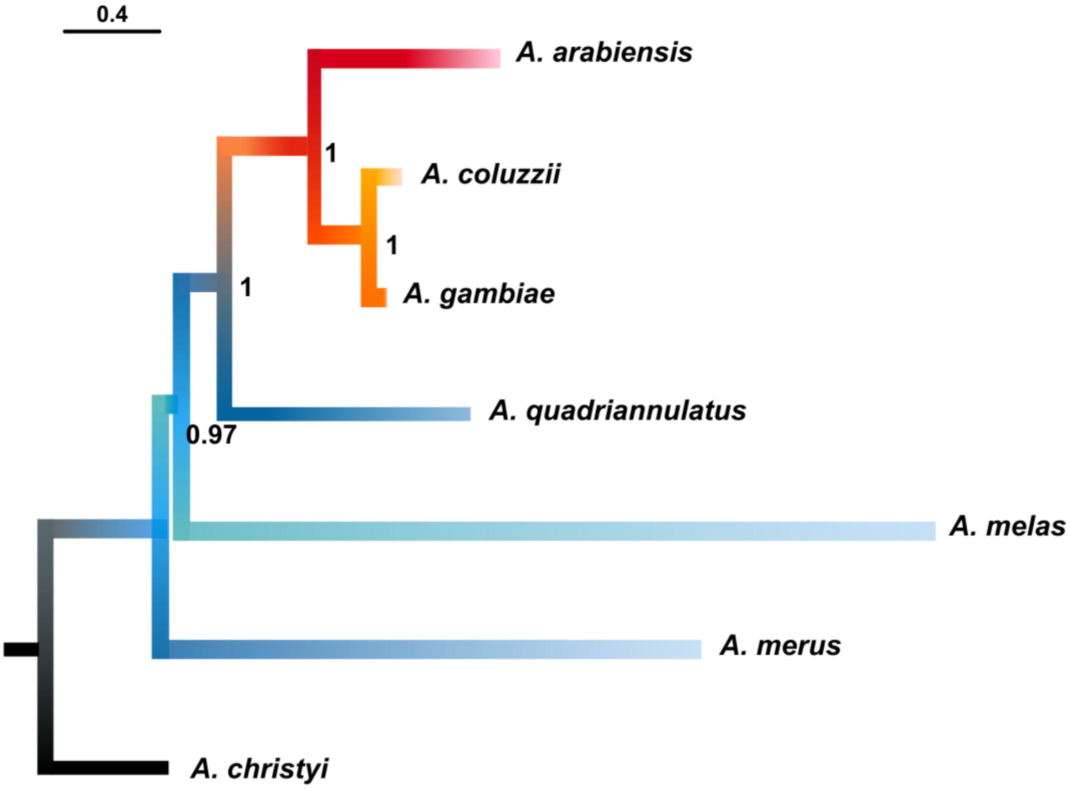

**Fig. S2.**
Coalescent-based species tree inferred from autosomal noncoding loci using ASTRAL-III. The species tree was summarized from individual locus trees inferred for 4,214 autosomal noncoding loci. Locus trees were inferred with IQ-TREE2 and rooted with *A. christyi* before analysis with ASTRAL. Node labels indicate local posterior probabilities, and the scale bar denotes branch length in coalescent units.

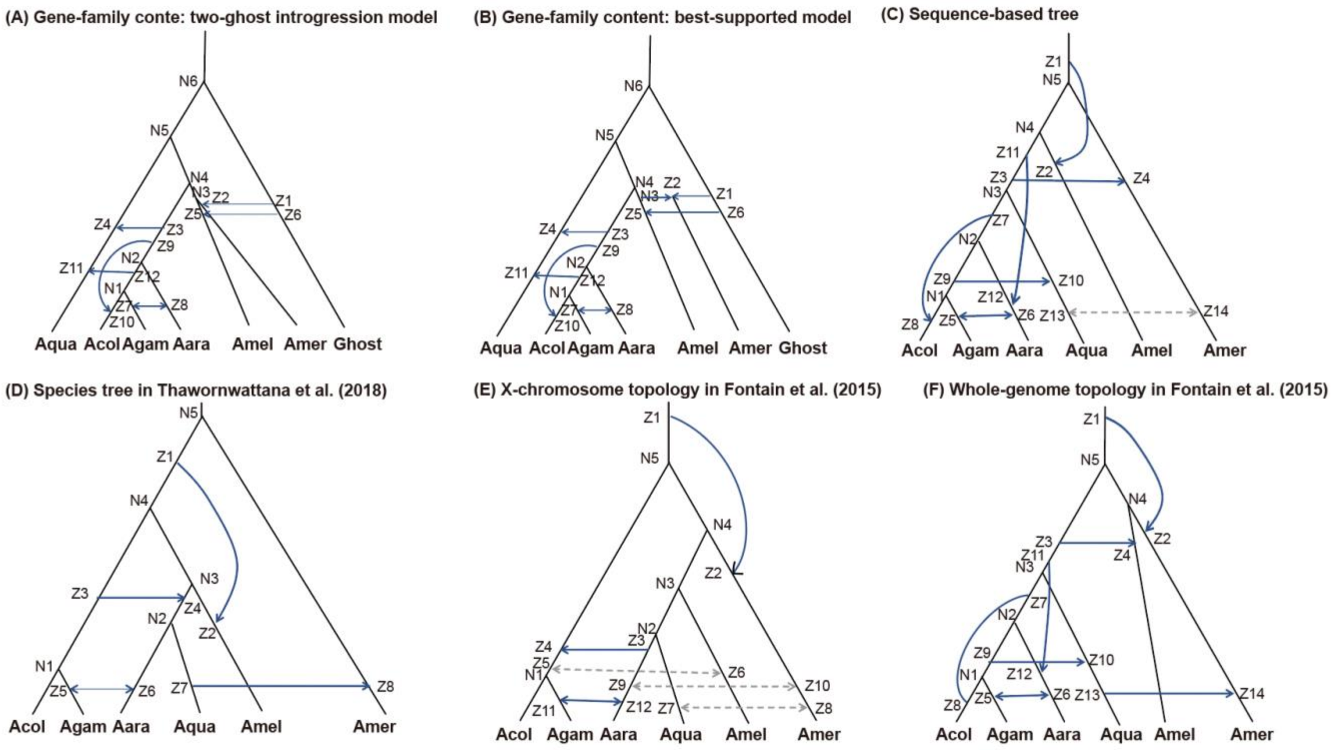

**Fig. S3.**

**D-BPP networks inferred under alternative species-tree hypotheses and competing ghost-ancestry models.** **(A)** Two-ghost introgression model inferred under the gene-family content species-tree topology, with separate post-divergence ghost-introgression events involving *A. merus* and *A. melas*. **(B)** Best-supported model under the same gene-family content topology, in which *A. merus* is modeled as a ghost-mediated hybrid lineage and *A. melas* retains a post-divergence ghost-introgression event. **(C)** Network inferred under the sequence-based ASTRAL species tree. **(D)** Network inferred under the species tree reported by Thawornwattana et al. (2018). **(E)** Network inferred under the X-chromosome topology reported by Fontaine et al. (2015). (F) Network inferred under the whole-genome tree reported by Fontaine et al. (2015). Panels A and B represent the two competing models specifically compared under the gene-family content topology; the best-supported model in panel B was subsequently used in the cross-topology marginal-likelihood comparison. Blue arrows indicate reticulation events retained in the corresponding D-BPP network, whereas dashed gray arrows indicate candidate events that were evaluated but not retained. Labels beginning with N denote divergence nodes, and labels beginning with Z denote reticulation-related nodes in the corresponding MSci models. Abbreviations: Acol, *A. coluzzii*; Agam, *A. gambiae*; Aara, *A. arabiensis*; Aqua, *A. quadriannulatus*; Amel, *A. melas*; Amer, *A. merus*.

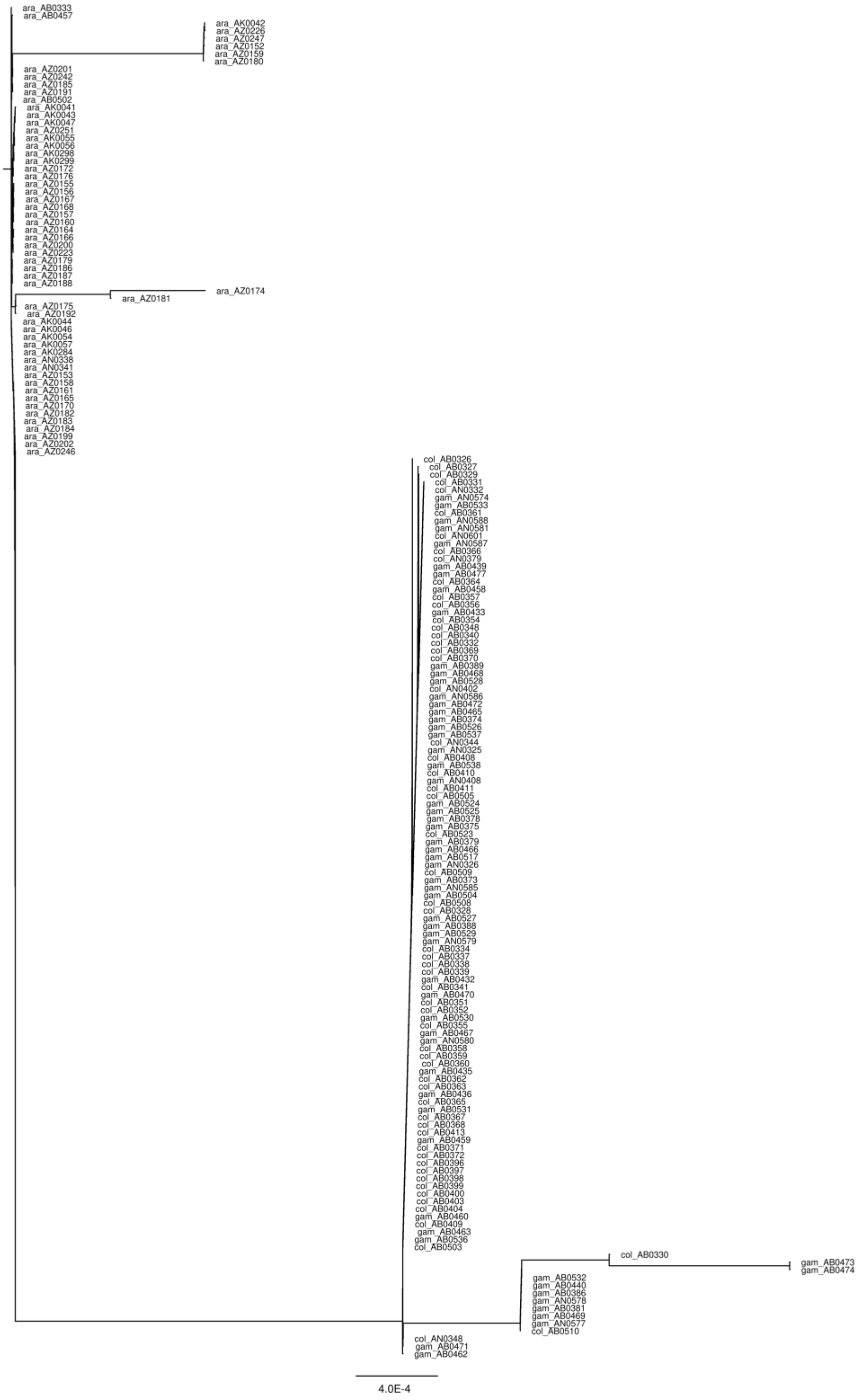

**Fig. S4.**
Individual-level BioNJ gene tree for OBP4 (AGAP010489) in the chromosome arm 3L candidate interval. The tree includes *A. gambiae* (gam), *A. coluzzii* (col), and *A. arabiensis* (ara).

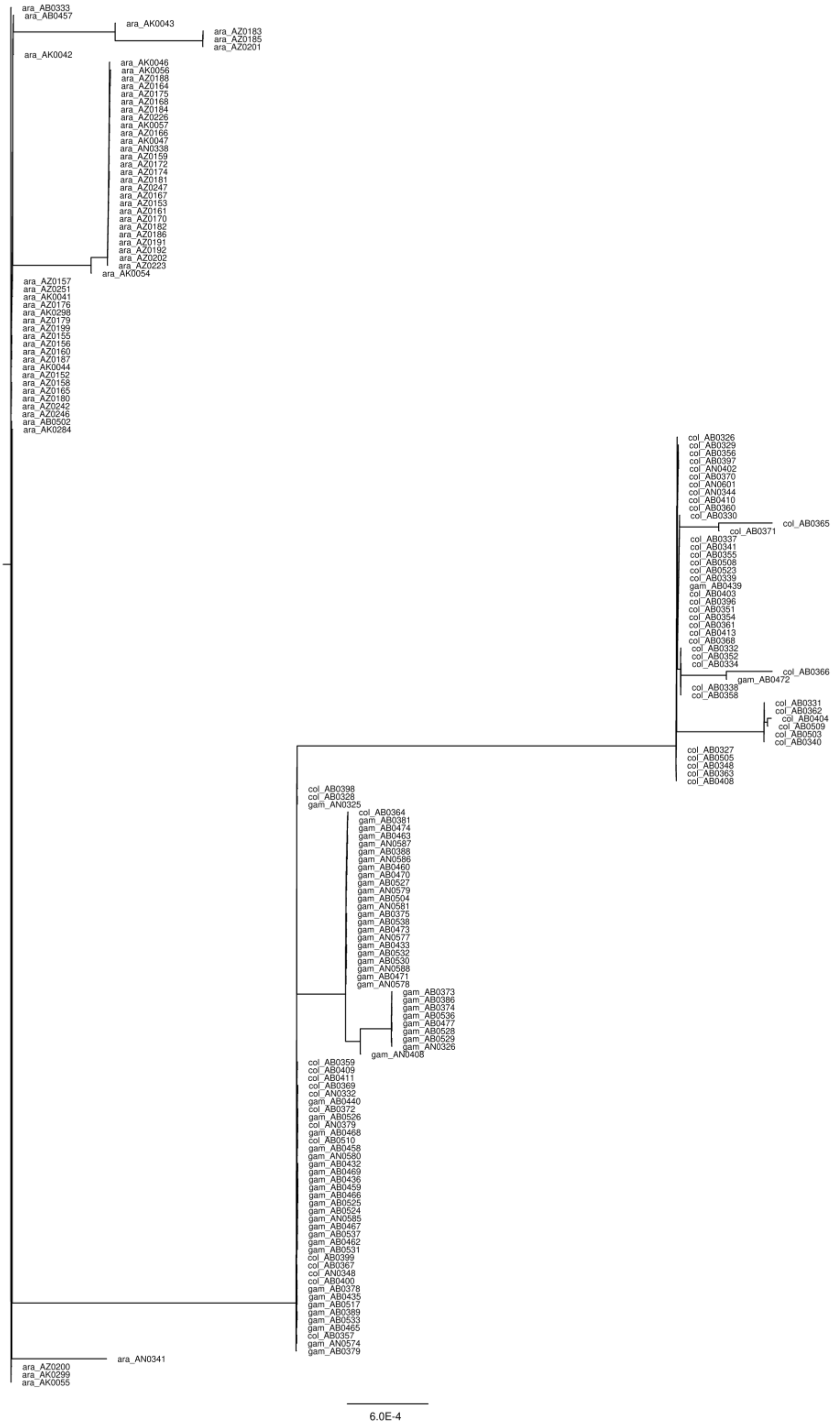

**Fig. S5.**
Individual-level BioNJ gene tree for Or43 (AGAP010504) in the chromosome arm 3L candidate interval. The tree includes *A. gambiae* (gam), *A. coluzzii* (col), and *A. arabiensis* (ara).

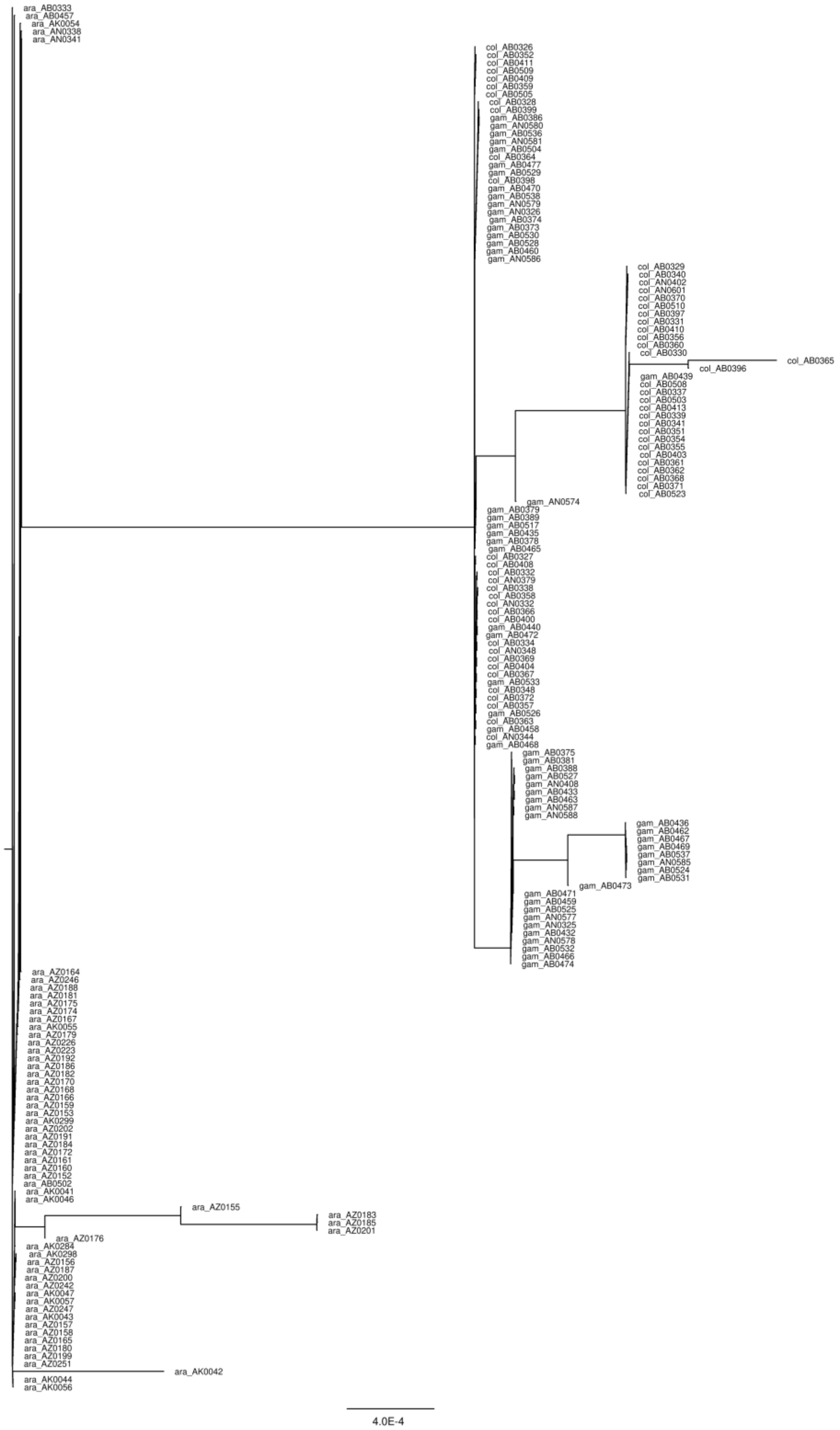

**Fig. S6.**
Individual-level BioNJ gene tree for Or44 (AGAP010505) in the chromosome arm 3L candidate interval. The tree includes *A. gambiae* (gam), *A. coluzzii* (col), and *A. arabiensis* (ara).

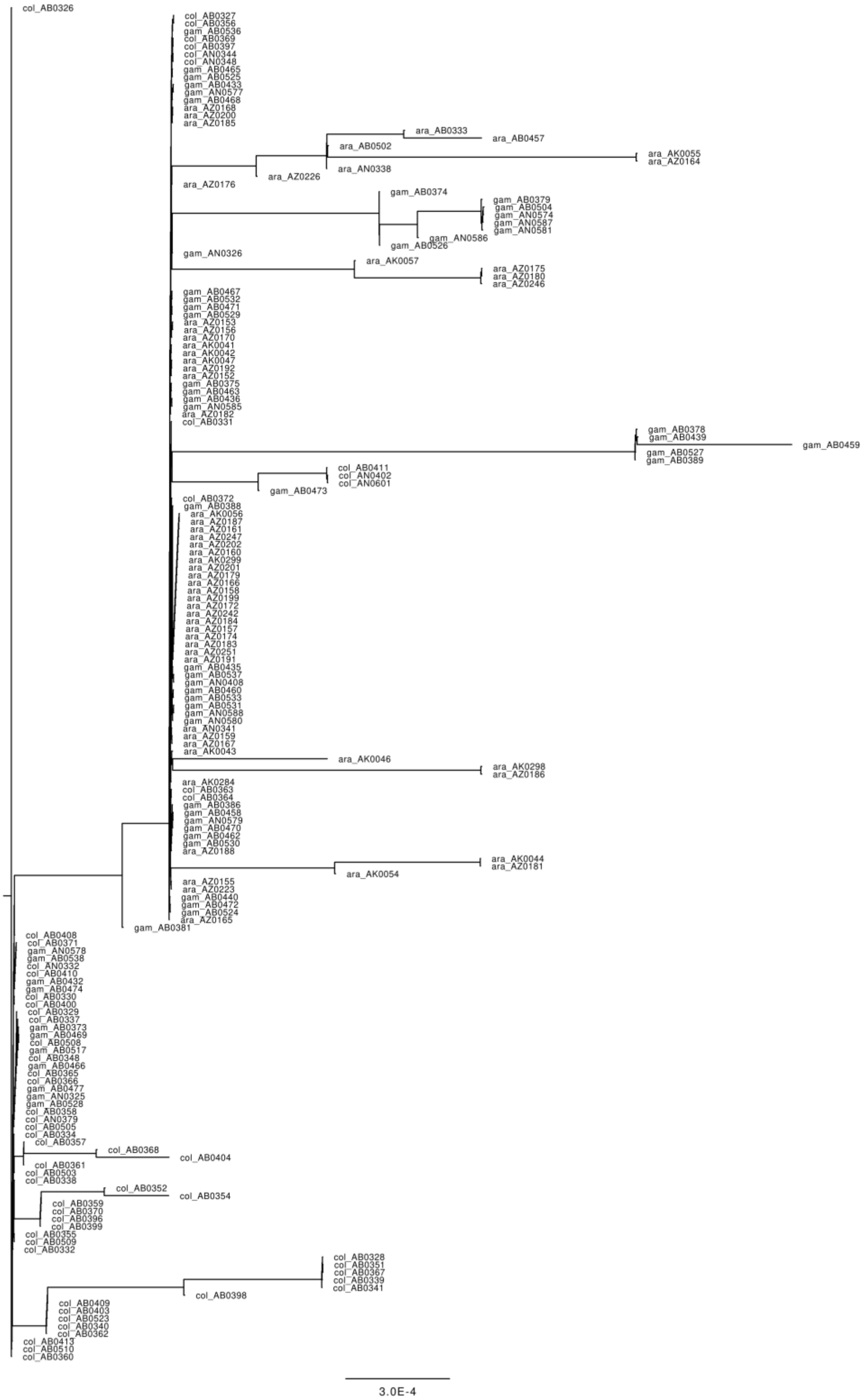

**Fig. S7.**
Individual-level BioNJ gene tree for AGAP010105 within the RR-2 cuticular-protein cluster in the chromosome arm 3R candidate interval. The tree includes *A. gambiae* (gam), *A. coluzzii* (col), and *A. arabiensis* (ara).

**Table S1. Gene-content phylogenetic trees inferred from matrix perturbation tests across three perturbation levels.**

| Perturbation level | Replicate | Inferred Newick tree |
|---|---|---|
| 0.001 | 1 | (Aepi:0.0308602,(Achr:0.3512197211,(Aqua:0.0721317660,((Amel:0.2352256291,Amer:0.0660155196)100:0.4540532023,(Aara:0.0476589128,(Acol:0.1068341824,Agam:0.0265741402)100:0.0600544617):0.0323103235)100:0.0287077665)90:0.1763839760)100:0.0308602); |
| 0.001 | 2 | (Aepi:0.0288065,(Achr:0.3509153699,(Aqua:0.0715381051,((Amel:0.2332300958,Amer:0.0623792876)100:0.4627880959,(Aara:0.0457371887,(Acol:0.1058237760,Agam:0.0261983759)100:0.0599598906):0.0317694407)100:0.0298412047)93:0.1778663495)100:0.0288065); |
| 0.001 | 3 | (Aepi:0.0301972,(Achr:0.3528344999,(Aqua:0.0727242723,((Amel:0.2353137119,Amer:0.0668855700)100:0.4609772918,(Aara:0.0466947610,(Acol:0.1060342737,Agam:0.0270387030)100:0.0610314827):0.0330506119)100:0.0285634029)91:0.1795992099)100:0.0301972); |
| 0.001 | 4 | (Aepi:0.0313212,(Achr:0.3503654506,(Aqua:0.0721117589,((Amel:0.2361265237,Amer:0.0665177871)100:0.4569047480,(Aara:0.0468840412,(Acol:0.1070246569,Agam:0.0261890785)100:0.0608886798):0.0326903429)100:0.0279711820)92:0.1767560883)100:0.0313212); |
| 0.001 | 5 | (Aepi:0.0309527,(Achr:0.3517869667,(Aqua:0.0723371732,((Amel:0.2371738485,Amer:0.0656737249)100:0.4610604014,(Aara:0.0474872516,(Acol:0.1064584826,Agam:0.0260309408)100:0.0605649463):0.0322514569)100:0.0289707126)90:0.1769553462)100:0.0309527); |
| 0.001 | 6 | (Aepi:0.0302065,(Achr:0.3520430055,(Aqua:0.0715450479,((Amel:0.2318163590,Amer:0.0661761113)100:0.4646542879,(Aara:0.0455167437,(Acol:0.1059284433,Agam:0.0275510023)100:0.0601570910):0.0341521349)100:0.0271215380)90:0.1781301868)100:0.0302065); |
| 0.001 | 7 | (Aepi:0.0308019,(Achr:0.3528276336,(Aqua:0.0724064618,((Amel:0.2361674154,Amer:0.0671772146)100:0.4594831653,(Aara:0.0472361714,(Acol:0.1066489258,Agam:0.0267020846)100:0.0612655261):0.0324433009)100:0.0283899605)91:0.1783017418)100:0.0308019); |
| 0.001 | 8 | (Aepi:0.0299179,(Achr:0.3551526190,(Aqua:0.0724758388,((Amel:0.2327567246,Amer:0.0675081526)100:0.4639896687,(Aara:0.0464837985,(Acol:0.1064394128,Agam:0.0260300083)100:0.0610642586):0.0330714029)100:0.0277906340)94:0.1811136088)100:0.0299179); |
| 0.001 | 9 | (Aepi:0.0294892,(Achr:0.3488438458,(Aqua:0.0708541978,((Amel:0.2310838319,Amer:0.0645561417)100:0.4570479349,(Aara:0.0465502686,(Acol:0.1053694097,Agam:0.0257340214)100:0.0597262805):0.0320518796)100:0.0284948289)93:0.1778181941)100:0.0294892); |
| 0.001 | 10 | (Aepi:0.0309421,(Achr:0.3501205844,(Aqua:0.0724222449,((Amel:0.2318271744,Amer:0.0682282702)100:0.4602435361,(Aara:0.0465886608,(Acol:0.1062276749,Agam:0.0252798049)100:0.0611728562):0.0339291117)100:0.0266797664)86:0.1773467249)100:0.0309421); |
| 0.005 | 1 | (Aepi:0.0316207,(Achr:0.3686580467,(Aqua:0.0777268866,((Amel:0.2442417628,Amer:0.0716536419)100:0.4881592790,(Aara:0.0517332802,(Acol:0.1129040987,Agam:0.0295295661)100:0.0633330990):0.0359438211)100:0.0255825544)83:0.1859518452)100:0.0316207); |
| 0.005 | 2 | (Aepi:0.0331955,(Achr:0.3704860446,(Aqua:0.0767935955,((Amel:0.2444549383,Amer:0.0696875153)100:0.4982441959,(Aara:0.0527054441,(Acol:0.1116364198,Agam:0.0297762821)100:0.0635653931):0.0347231438)100:0.0287913427)90:0.1830322876)100:0.0331955); |
| 0.005 | 3 | (Aepi:0.0343151,(Achr:0.3654704368,(Aqua:0.0777130582,((Amel:0.2465662438,Amer:0.0690593964)100:0.4826044054,(Aara:0.0511971752,(Acol:0.1125088712,Agam:0.0309455973)100:0.0625642420):0.0333897000)100:0.0279213136)90:0.1849684161)100:0.0343151); |
| 0.005 | 4 | (Aepi:0.0314435,(Achr:0.3797638649,(Aqua:0.0755004421,((Amel:0.2557876403,Amer:0.0665449438)100:0.4949184916,(Aara:0.0526233414,(Acol:0.1127567157,Agam:0.0298903595)100:0.0622107900):0.0334085408)100:0.0305949106)95:0.1885240765)100:0.0314435); |
| 0.005 | 5 | (Aepi:0.0323587,(Achr:0.3713590298,(Aqua:0.0770305995,((Amel:0.2488710460,Amer:0.0631690644)100:0.4878772205,(Aara:0.0510842659,(Acol:0.1111589724,Agam:0.0287415779)100:0.0638360366):0.0313175723)99:0.0295521348)96:0.1819456311)100:0.0323587); |
| 0.005 | 6 | (Aepi:0.0333355,(Achr:0.3645585138,(Aqua:0.0775850627,((Amel:0.2514724939,Amer:0.0721335970)100:0.4778723623,(Aara:0.0516422950,(Acol:0.1123271786,Agam:0.0312490051)100:0.0632761191):0.0341701456)100:0.0281357043)91:0.1835343790)100:0.0333355); |
| 0.005 | 7 | (Aepi:0.0305833,(Achr:0.3652399863,(Aqua:0.0733405807,((Amel:0.2466882274,Amer:0.0664043127)100:0.4817346479,(Aara:0.0478927149,(Acol:0.1115761433,Agam:0.0295176904)100:0.0620439180):0.0355282498)100:0.0298046533)92:0.1877126458)100:0.0305833); |
| 0.005 | 8 | (Aepi:0.031156,(Achr:0.3733658060,(Aqua:0.0784425530,((Amel:0.2529218225,Amer:0.0682709684)100:0.4925738134,(Aara:0.0517855635,(Acol:0.1149457086,Agam:0.0289802175)100:0.0626492988):0.0322896943)99:0.0305497537)95:0.1885295510)100:0.031156); |
| 0.005 | 9 | (Aepi:0.0344264,(Achr:0.3639624051,(Aqua:0.0771360509,((Amel:0.2436644902,Amer:0.0724913074)100:0.4781158878,(Aara:0.0496570113,(Acol:0.1104617785,Agam:0.0290424472)100:0.0632172900):0.0328613507)100:0.0275364562)89:0.1806667207)100:0.0344264); |
| 0.005 | 10 | (Aepi:0.0318198,(Achr:0.3684417839,(Aqua:0.0794125281,((Amel:0.2394572101,Amer:0.0654211725)100:0.4940501068,(Aara:0.0506653940,(Acol:0.1116622260,Agam:0.0314153790)100:0.0619265685):0.0363888756)100:0.0273453982)89:0.1828847046)100:0.0318198); |
| 0.01 | 1 | (Aepi:0.0326388,(Achr:0.3935215000,(Aqua:0.0836139649,((Amel:0.2695378753,Amer:0.0620788725)100:0.5347799580,(Aara:0.0565020641,(Acol:0.1184636670,Agam:0.0346953609)100:0.0643127123):0.0344833364)99:0.0290782843)94:0.1977248258)100:0.0326388); |
| 0.01 | 2 | (Aepi:0.0318888,(Achr:0.3938716847,(Aqua:0.0820351536,((Amel:0.2570962020,Amer:0.0696967668)100:0.5226351844,(Aara:0.0563443419,(Acol:0.1177058581,Agam:0.0336404139)100:0.0664961995):0.0347266320)99:0.0317633320)95:0.1988949879)100:0.0318888); |
| 0.01 | 3 | (Aepi:0.0320934,(Achr:0.4061461446,(Aqua:0.0813248046,((Amel:0.2633679007,Amer:0.0758414360)100:0.5296869982,(Aara:0.0576317618,(Acol:0.1202427126,Agam:0.0321310866)100:0.0688431286):0.0330608129)99:0.0354312507)98:0.1999859811)100:0.0320934); |
| 0.01 | 4 | (Aepi:0.0332169,(Achr:0.3874653274,(Aqua:0.0811557780,((Amel:0.2626722002,Amer:0.0806654499)100:0.5039136603,(Aara:0.0559695599,(Acol:0.1210563102,Agam:0.0371603714)100:0.0656346867):0.0336846063)99:0.0308945073)97:0.1928196432)100:0.0332169); |
| 0.01 | 5 | (Aepi:0.0277931,(Achr:0.4101264594,(Aqua:0.0841262708,((Amel:0.2676217272,Amer:0.0773716717)100:0.5405981854,(Aara:0.0545889212,(Acol:0.1200224677,Agam:0.0326423659)100:0.0649968469):0.0424789679)100:0.0234632896)88:0.2084993069) |

| Perturbation level | Replicate | Inferred Newick tree |
|---|---|---|
| | | 100:0.0277931); |
| 0.01 | 6 | (Aepi:0.0347458,(Achr:0.3817965698,(Aqua:0.0807815448,((Amel:0.2490864228,Amer:0.0739552139)100:0.4988980633,(Aara: 0.0538228846,(Acol:0.1197612551,Agam:0.0345733898)100:0.0624577073):0.0324207827)99:0.0337521740)95:0.1864363125)1 00:0.0347458); |
| 0.01 | 7 | (Aepi:0.0310465,(Achr:0.3886530669,(Aqua:0.0829155857,((Amel:0.2650136317,Amer:0.0624699049)100:0.5341507950,(Aara: 0.0571840828,(Acol:0.1167694729,Agam:0.0347479482)100:0.0632807115):0.0335765665)99:0.0292081096)92:0.2007268237)1 00:0.0310465); |
| 0.01 | 8 | (Aepi:0.030631,(Achr:0.3816143944,(Aqua:0.0813857401,((Amel:0.2646414871,Amer:0.0647605290)100:0.5125267470,(Aara:0. 0559358989,(Acol:0.1163315569,Agam:0.0355168650)100:0.0646646338):0.0331537597)99:0.0333101903)95:0.1940373369)10 0:0.030631); |
| 0.01 | 9 | (Aepi:0.0307416,(Achr:0.3966560117,(Aqua:0.0797820067,((Amel:0.2703634569,Amer:0.0646410719)100:0.5431853562,(Aara: 0.0524008923,(Acol:0.1224822219,Agam:0.0338784381)100:0.0692992938):0.0349348355)99:0.0323901807)98:0.2039105931)1 00:0.0307416); |
| 0.01 | 10 | (Aepi:0.0371432,(Achr:0.3823855093,(Aqua:0.0823233230,((Amel:0.2649244444,Amer:0.0760792223)100:0.5114308053,(Aara: 0.0561182273,(Acol:0.1195592588,Agam:0.0332066719)100:0.0651441584):0.0342295367)100:0.0305162810)95:0.1912980300) 100:0.0371432); |

### Table S2. Savage–Dickey density-ratio tests in simulated hybrid-speciation scenarios.

Bayes factors (Bε) were calculated to test equality between the relevant species-divergence and reticulation time parameters under three null intervals. Simulations include standard hybrid speciation with two sampled parental lineages and ghost-mediated hybrid speciation with one unsampled parental lineage.

| Scenario | Model / parental sampling | No. loci | Replicate | Bε (ε = $10^{-4}$) | Bε (ε = $10^{-5}$) | Bε (ε = $10^{-6}$) |
|---|---|---|---|---|---|---|
| Hybrid speciation | Two sampled parental lineages | 5000 | 1 | 0.0202 | 0.0099 | 0.0073 |
| Hybrid speciation | Two sampled parental lineages | 5000 | 2 | 0.0152 | 0.0076 | 0.0068 |
| Hybrid speciation | Two sampled parental lineages | 5000 | 3 | 0.0669 | 0.1229 | 0.0133 |
| Hybrid speciation | Two sampled parental lineages | 5000 | 4 | 0.0457 | 0.0967 | 0.1866 |
| Hybrid speciation | Two sampled parental lineages | 5000 | 5 | 0.0541 | 0.0524 | 0.0056 |
| Hybrid speciation | Two sampled parental lineages | 5000 | 6 | 0.0236 | 0.0171 | 0.0102 |
| Hybrid speciation | Two sampled parental lineages | 5000 | 7 | 0.025 | 0.0475 | 0.0165 |
| Hybrid speciation | Two sampled parental lineages | 5000 | 8 | 0.0199 | 0.0163 | 0.007 |
| Hybrid speciation | Two sampled parental lineages | 5000 | 9 | 0.0134 | 0.0088 | 0.0023 |
| Hybrid speciation | Two sampled parental lineages | 5000 | 10 | 0.0347 | 0.0476 | 0.0464 |
| Ghost-mediated hybrid speciation | One unsampled parental lineage (C = ghost) | 5000 | 1 | 0.0588 | 0.0426 | 0.0292 |
| Ghost-mediated hybrid speciation | One unsampled parental lineage (C = ghost) | 5000 | 2 | 0.0438 | 0.0335 | 0.0111 |
| Ghost-mediated hybrid speciation | One unsampled parental lineage (C = ghost) | 5000 | 3 | 0.2893 | 0.4255 | 0.2941 |
| Ghost-mediated hybrid speciation | One unsampled parental lineage (C = ghost) | 5000 | 4 | 0.155 | 0.1848 | 0.0309 |
| Ghost-mediated hybrid speciation | One unsampled parental lineage (C = ghost) | 5000 | 5 | 0.0281 | 0.0212 | 0.014 |
| Ghost-mediated hybrid speciation | One unsampled parental lineage (C = ghost) | 5000 | 6 | 0.0919 | 0.0903 | 0.0569 |
| Ghost-mediated hybrid speciation | One unsampled parental lineage (C = ghost) | 5000 | 7 | 0.0353 | 0.0322 | 0.0644 |
| Ghost-mediated hybrid speciation | One unsampled parental lineage (C = ghost) | 5000 | 8 | 0.326 | 0.508 | 0.1618 |
| Ghost-mediated hybrid speciation | One unsampled parental lineage (C = ghost) | 5000 | 9 | 0.0618 | 0.0473 | 0.026 |
| Ghost-mediated hybrid speciation | One unsampled parental lineage (C = ghost) | 5000 | 10 | 0.036 | 0.0302 | 0.0362 |

### Table S3. D-statistic results based on the gene-family content species-tree topology

| P1 | P2 | P3 | D | Z-score | p-value | f4-ratio | BBAA | ABBA | BABA | Dp | adjusted_p_value |
|---|---|---|---|---|---|---|---|---|---|---|---|
| Amer | Acol | Aqua | 0.363807 | 35.0757 | 2.3e-16 | 0.279899 | 2451 | 5799.25 | 2705.25 | 0.282415 | 4.6e-15 |
| Amer | Agam | Aqua | 0.363626 | 31.9143 | 2.3e-16 | 0.279427 | 2447.5 | 5792 | 2703 | 0.282294 | 4.6e-15 |
| Amer | Aara | Aqua | 0.334179 | 28.8952 | 2.3e-16 | 0.254503 | 2496.12 | 5584.38 | 2786.88 | 0.257422 | 4.6e-15 |
| Amel | Agam | Aqua | 0.335109 | 26.2151 | 2.3e-16 | 0.241653 | 2646.12 | 5042.88 | 2511.38 | 0.248177 | 4.6e-15 |
| Amel | Acol | Aqua | 0.335425 | 25.9988 | 2.3e-16 | 0.241215 | 2661.12 | 5028.38 | 2502.38 | 0.247844 | 4.6e-15 |
| Amel | Aara | Aqua | 0.295576 | 24.6999 | 2.3e-16 | 0.213845 | 2683.88 | 4879.62 | 2653.12 | 0.217929 | 4.6e-15 |
| Amer | Amel | Acol | 0.111683 | 10.584 | 2.3e-16 | 0.0930085 | 2990.75 | 4750.5 | 3796 | 0.082732 | 4.6e-15 |
| Amer | Amel | Agam | 0.107624 | 10.2884 | 2.3e-16 | 0.0917003 | 2984 | 4703.25 | 3789.25 | 0.079641 | 4.6e-15 |
| Amer | Amel | Aara | 0.101584 | 8.8372 | 2.3e-16 | 0.0779344 | 3064.25 | 4676.5 | 3814 | 0.0746446 | 4.6e-15 |
| Amer | Amel | Aqua | 0.0624855 | 5.87554 | 4.21476e-09 | 0.0491308 | 2984 | 4565.5 | 4028.5 | 0.0463811 | 8.42952e-08 |
| Acol | Agam | Aara | 0.0680161 | 5.44107 | 5.29615e-08 | 0.0394613 | 3521.75 | 1886.25 | 1646 | 0.0340587 | 1.05923e-06 |
| Aara | Agam | Aqua | 0.0684413 | 4.84418 | 1.27137e-06 | 0.0344088 | 4328.88 | 2222.62 | 1937.88 | 0.0335407 | 2.54274e-05 |
| Aara | Acol | Aqua | 0.0633936 | 4.83405 | 1.33786e-06 | 0.0333605 | 4191.12 | 2314.88 | 2038.88 | 0.0323 | 2.67572e-05 |

### Table S4. Summary of Bayesian tests for gene flow using BPP under different species-tree topologies

| Topology | Gene-flow event | B10 (ε=0.01) |
|---|---|---|
| Gene-family content | Z1→Z2 | ∞ |
| | Z3→Z4 | ∞ |
| | Z6→Z5 | ∞ |
| | Z7→Z8 | ∞ |
| | Z8→Z7 | ∞ |
| | Z9→Z10 | ∞ |
| | Z12→Z11 | ∞ |
| Sequence-based tree | Z1→Z2 | ∞ |
| | Z3→Z4 | ∞ |
| | Z5→Z6 | ∞ |
| | Z6→Z5 | ∞ |
| | Z7→Z8 | ∞ |
| | Z9→Z10 | ∞ |
| | Z10→Z9 | 0.01 |
| | Z11→Z12 | ∞ |
| | Z13→Z14 | 0.01 |
| | Z14→Z13 | 0.01 |
| Species tree in Thawornwattana et al. (2018) | Z1→Z2 | ∞ |
| | Z3→Z4 | ∞ |
| | Z5→Z6 | ∞ |
| | Z6→Z5 | ∞ |
| | Z7→Z8 | ∞ |
| | Z8→Z7 | 0.01 |
| Whole-genome tree in Fontaine et al. (2015) | Z1→Z2 | ∞ |
| | Z3→Z4 | ∞ |
| | Z5→Z6 | ∞ |
| | Z6→Z5 | ∞ |
| | Z7→Z8 | ∞ |
| | Z9→Z10 | ∞ |
| | Z10→Z9 | 1.71 |
| | Z11→Z12 | ∞ |
| | Z13→Z14 | ∞ |
| | Z14→Z13 | 0.01 |
| X-chromosome topology in Fontaine et al. (2015) | Z1→Z2 | ∞ |
| | Z3→Z4 | ∞ |
| | Z4→Z3 | 0.24 |
| | Z5→Z6 | 1.31 |
| | Z6→Z5 | 0.01 |
| | Z7→Z8 | 0.8 |
| | Z8→Z7 | 0.05 |
| | Z9→Z10 | 0.08 |
| | Z10→Z9 | 0.01 |
| | Z11→Z12 | ∞ |
| | Z12→Z11 | ∞ |

The corresponding positions of all node labels in the MSci network are shown in Fig. S3.

### Table S5. D-statistic results based on the sequence-based species-tree topology

| P1 | P2 | P3 | D | Z-score | p-value | f4-ratio | BBAA | ABBA | BABA | Dp | adjusted_p_value |
|---|---|---|---|---|---|---|---|---|---|---|---|
| Amel | Aqua | Amer | 0.148948 | 16.9151 | 2.3e-16 | 0.06921 | 4565.5 | 4028.5 | 2984 | 0.0902142 | 4.6e-15 |
| Amel | Agam | Amer | 0.118887 | 12.1428 | 2.3e-16 | 0.0530739 | 4703.25 | 3789.25 | 2984 | 0.0701651 | 4.6e-15 |
| Amel | Acol | Amer | 0.11865 | 11.3131 | 2.3e-16 | 0.0530538 | 4750.5 | 3796 | 2990.75 | 0.0697957 | 4.6e-15 |
| Amel | Aara | Amer | 0.109003 | 12.858 | 2.3e-16 | 0.049714 | 4676.5 | 3814 | 3064.25 | 0.0648867 | 4.6e-15 |
| Acol | Agam | Aara | 0.0680161 | 5.44107 | 5.29615e-08 | 0.0394726 | 3521.75 | 1886.25 | 1646 | 0.0340587 | 1.05923e-06 |
| Aara | Agam | Aqua | 0.0684413 | 4.84418 | 1.27137e-06 | 0.0342073 | 4328.88 | 2222.62 | 1937.88 | 0.0335407 | 2.54274e-05 |
| Aara | Acol | Aqua | 0.0633936 | 4.83405 | 1.33786e-06 | 0.0331611 | 4191.12 | 2314.88 | 2038.88 | 0.0323 | 2.67572e-05 |
| Aara | Aqua | Amer | 0.055035 | 4.84624 | 1.25822e-06 | 0.0201959 | 5584.38 | 2786.88 | 2496.12 | 0.0267553 | 2.51644e-05 |
| Agam | Aqua | Amer | 0.0496068 | 3.57028 | 0.000356595 | 0.0176936 | 5792 | 2703 | 2447.5 | 0.0233493 | 0.0071319 |
| Acol | Aqua | Amer | 0.0493091 | 4.2965 | 1.7352e-05 | 0.017611 | 5799.25 | 2705.25 | 2451 | 0.0232075 | 0.00034704 |

### Table S6. D-statistic results based on the species topology in Thawornwattana et al. (2018)

| P1 | P2 | P3 | D | Z-score | p-value | f4-ratio | BBAA | ABBA | BABA | Dp | adjusted_p_value |
|---|---|---|---|---|---|---|---|---|---|---|---|
| Amel | Aara | Agam | 0.53127 | 43.6191 | 2.3e-16 | 0.491376 | 1868.5 | 6445.5 | 1973 | 0.434772 | 4.6e-15 |
| Amel | Aara | Acol | 0.506913 | 37.7728 | 2.3e-16 | 0.455973 | 1918.75 | 6267.25 | 2050.75 | 0.411898 | 4.6e-15 |
| Aqua | Aara | Agam | 0.321491 | 17.374 | 2.3e-16 | 0.311506 | 1937.88 | 4328.88 | 2222.62 | 0.248105 | 4.6e-15 |
| Amel | Aqua | Agam | 0.311712 | 21.6897 | 2.3e-16 | 0.263206 | 2511.38 | 5042.88 | 2646.12 | 0.234968 | 4.6e-15 |
| Amel | Aqua | Acol | 0.307855 | 21.9079 | 2.3e-16 | 0.256168 | 2502.38 | 5028.38 | 2661.12 | 0.232269 | 4.6e-15 |
| Aqua | Aara | Acol | 0.288388 | 17.3999 | 2.3e-16 | 0.270821 | 2038.88 | 4191.12 | 2314.88 | 0.219575 | 4.6e-15 |
| Amel | Aqua | Amer | 0.148948 | 16.9151 | 2.3e-16 | 0.0692203 | 4565.5 | 4028.5 | 2984 | 0.0902142 | 4.6e-15 |
| Amel | Agam | Amer | 0.118887 | 12.1428 | 2.3e-16 | 0.05308 | 4703.25 | 3789.25 | 2984 | 0.0701651 | 4.6e-15 |
| Amel | Acol | Amer | 0.11865 | 11.3131 | 2.3e-16 | 0.0530643 | 4750.5 | 3796 | 2990.75 | 0.0697957 | 4.6e-15 |
| Amel | Aara | Amer | 0.109003 | 12.858 | 2.3e-16 | 0.0497157 | 4676.5 | 3814 | 3064.25 | 0.0648867 | 4.6e-15 |
| Acol | Agam | Aara | 0.0680161 | 5.44107 | 5.29615e-08 | 0.0396142 | 3521.75 | 1886.25 | 1646 | 0.0340587 | 1.05923e-06 |
| Aara | Aqua | Amer | 0.055035 | 4.84624 | 1.25822e-06 | 0.0201931 | 5584.38 | 2786.88 | 2496.12 | 0.0267553 | 2.51644e-05 |
| Agam | Aqua | Amer | 0.0496068 | 3.57028 | 0.000356595 | 0.0176869 | 5792 | 2703 | 2447.5 | 0.0233493 | 0.0071319 |
| Acol | Aqua | Amer | 0.0493091 | 4.2965 | 1.7352e-05 | 0.017607 | 5799.25 | 2705.25 | 2451 | 0.0232075 | 0.00034704 |

### Table S7. D-statistic results based on the X-chromosome topology in Fontaine et al. (2015)

| P1 | P2 | P3 | D | Z-score | p-value | f4-ratio | BBAA | ABBA | BABA | Dp | adjusted_p_value |
|---|---|---|---|---|---|---|---|---|---|---|---|
| Amer | Aara | Agam | 0.587096 | 44.5728 | 2.3e-16 | 0.542553 | 1863.62 | 7345.88 | 1911.12 | 0.48871 | 4.6e-15 |
| Amer | Aara | Acol | 0.565077 | 41.7091 | 2.3e-16 | 0.505657 | 1963.88 | 7225.38 | 2007.88 | 0.465967 | 4.6e-15 |
| Amel | Aara | Agam | 0.53127 | 43.6191 | 2.3e-16 | 0.492254 | 1868.5 | 6445.5 | 1973 | 0.434772 | 4.6e-15 |
| Amel | Aara | Acol | 0.506913 | 37.7728 | 2.3e-16 | 0.452815 | 1918.75 | 6267.25 | 2050.75 | 0.411898 | 4.6e-15 |
| Amer | Aqua | Agam | 0.405911 | 28.4597 | 2.3e-16 | 0.334291 | 2703 | 5792 | 2447.5 | 0.305643 | 4.6e-15 |
| Amer | Aqua | Acol | 0.405836 | 30.9141 | 2.3e-16 | 0.325199 | 2705.25 | 5799.25 | 2451 | 0.305623 | 4.6e-15 |
| Aqua | Aara | Agam | 0.321491 | 17.374 | 2.3e-16 | 0.313558 | 1937.88 | 4328.88 | 2222.62 | 0.248105 | 4.6e-15 |
| Amel | Aqua | Agam | 0.311712 | 21.6897 | 2.3e-16 | 0.263843 | 2511.38 | 5042.88 | 2646.12 | 0.234968 | 4.6e-15 |
| Amel | Aqua | Acol | 0.307855 | 21.9079 | 2.3e-16 | 0.254345 | 2502.38 | 5028.38 | 2661.12 | 0.232269 | 4.6e-15 |
| Aqua | Aara | Acol | 0.288388 | 17.3999 | 2.3e-16 | 0.267692 | 2038.88 | 4191.12 | 2314.88 | 0.219575 | 4.6e-15 |
| Amel | Aqua | Amer | 0.148948 | 16.9151 | 2.3e-16 | 0.0691436 | 4565.5 | 4028.5 | 2984 | 0.0902142 | 4.6e-15 |
| Amer | Amel | Acol | 0.111683 | 10.584 | 2.3e-16 | 0.0933679 | 2990.75 | 4750.5 | 3796 | 0.082732 | 4.6e-15 |
| Amer | Amel | Agam | 0.107624 | 10.2884 | 2.3e-16 | 0.0920188 | 2984 | 4703.25 | 3789.25 | 0.079641 | 4.6e-15 |
| Amel | Aara | Amer | 0.109003 | 12.858 | 2.3e-16 | 0.049686 | 4676.5 | 3814 | 3064.25 | 0.0648867 | 4.6e-15 |
| Acol | Agam | Aara | 0.0680161 | 5.44107 | 5.29615e-08 | 0.0396403 | 3521.75 | 1886.25 | 1646 | 0.0340587 | 1.05923e-06 |
| Aara | Aqua | Amer | 0.055035 | 4.84624 | 1.25822e-06 | 0.0201693 | 5584.38 | 2786.88 | 2496.12 | 0.0267553 | 2.51644e-05 |

### Table S8. D-statistic results based on the whole-genome topology in Fontaine et al. (2015)

| P1 | P2 | P3 | D | Z-score | p-value | f4-ratio | BBAA | ABBA | BABA | Dp | adjusted_p_value |
|---|---|---|---|---|---|---|---|---|---|---|---|
| Amer | Amel | Acol | 0.111683 | 10.584 | 2.3e-16 | 0.0935303 | 2990.75 | 4750.5 | 3796 | 0.082732 | 4.6e-15 |
| Amer | Amel | Agam | 0.107624 | 10.2884 | 2.3e-16 | 0.0915832 | 2984 | 4703.25 | 3789.25 | 0.079641 | 4.6e-15 |
| Amer | Amel | Aara | 0.101584 | 8.8372 | 2.3e-16 | 0.0778658 | 3064.25 | 4676.5 | 3814 | 0.0746446 | 4.6e-15 |
| Amer | Amel | Aqua | 0.0624855 | 5.87554 | 4.21476e-09 | 0.0493271 | 2984 | 4565.5 | 4028.5 | 0.0463811 | 8.42952e-08 |
| Acol | Agam | Aara | 0.0680161 | 5.44107 | 5.29615e-08 | 0.039471 | 3521.75 | 1886.25 | 1646 | 0.0340587 | 1.05923e-06 |
| Aara | Agam | Aqua | 0.0684413 | 4.84418 | 1.27137e-06 | 0.0345246 | 4328.88 | 2222.62 | 1937.88 | 0.0335407 | 2.54274e-05 |
| Aara | Acol | Aqua | 0.0633936 | 4.83405 | 1.33786e-06 | 0.0334667 | 4191.12 | 2314.88 | 2038.88 | 0.0323 | 2.67572e-05 |
| Aara | Aqua | Amer | 0.055035 | 4.84624 | 1.25822e-06 | 0.0202638 | 5584.38 | 2786.88 | 2496.12 | 0.0267553 | 2.51644e-05 |
| Agam | Aqua | Amer | 0.0496068 | 3.57028 | 0.000356595 | 0.0177477 | 5792 | 2703 | 2447.5 | 0.0233493 | 0.0071319 |
| Acol | Aqua | Amer | 0.0493091 | 4.2965 | 1.7352e-05 | 0.0176682 | 5799.25 | 2705.25 | 2451 | 0.0232075 | 0.00034704 |

**Table S9. Posterior parameter estimates for the two-ghost introgression model under the gene-family content species tree**

| node | τ | Time (Ma) | 95% HPD of τ | θ | 95% HPD of θ |
|---|---|---|---|---|---|
| N6 | 0.020975 | 0.6810 | 0.020452-0.021504 | 0.016010 | 0.015039-0.016984 |
| N5 | 0.018134 | 0.5888 | 0.017291-0.018965 | 0.003794 | 0.002768-0.004794 |
| N4 | 0.011886 | 0.3859 | 0.011664-0.012109 | 0.011498 | 0.010446-0.012593 |
| N3 | 0.011872 | 0.3855 | 0.011644-0.012095 | 0.013083 | 0.001219-0.028347 |
| Z1 | 0.011832 | 0.3842 | 0.011591-0.012062 | 0.190327 | 0.147622-0.234827 |
| Z2 | 0.011832 | 0.3842 | 0.011591-0.012062 | 0.009741 | 0.000103-0.023499 |
| Z5 | 0.011231 | 0.3646 | 0.010212-0.011972 | 0.009578 | 0.000157-0.023255 |
| Z4 | 0.009650 | 0.3133 | 0.009376-0.009921 | 0.007640 | 0.001549-0.015496 |
| Z3 | 0.009650 | 0.3133 | 0.009376-0.009921 | 0.014061 | 0.012257-0.015898 |
| Z9 | 0.009532 | 0.3095 | 0.009168-0.009868 | 0.010561 | 0.001111-0.022623 |
| N2 | 0.006727 | 0.2184 | 0.006477-0.006964 | 0.012595 | 0.011298-0.013896 |
| Z11 | 0.003568 | 0.1158 | 0.003125-0.003970 | 0.004894 | 0.004234-0.005586 |
| Z12 | 0.003568 | 0.1158 | 0.003125-0.003970 | 0.014478 | 0.013171-0.015770 |
| N1 | 0.002705 | 0.0878 | 0.002474-0.002949 | 0.017878 | 0.010198-0.024505 |
| Z7 | 0.002693 | 0.0874 | 0.002464-0.002940 | 0.012829 | 0.001200-0.027912 |
| Z8 | 0.002693 | 0.0874 | 0.002464-0.002940 | 0.004926 | 0.004368-0.005480 |
| Z10 | 0.002313 | 0.0751 | 0.001903-0.002678 | 0.021390 | 0.007948-0.037491 |

The corresponding positions of all node labels in the MSci network are shown in Fig. S3A.

**Table S10. Log marginal likelihoods of the two-ghost and hybrid-ghost MSci network models based on the gene-family content species tree.**

| Dataset | Partition | Two-ghost introgression model | hy+ghost model |
|---|---|---|---|
| odd | LNK1 | -704350.93 | -704142.34 |
| | LNK2 | -687139.01 | -686875.47 |
| even | LNK3 | -701829.37 | -701702.11 |
| | LNK4 | -660239.85 | -660138.73 |

**Table S11. Log marginal likelihoods of MSci networks inferred under alternative species-tree hypotheses and previously published network scenarios.**

| Dataset | Partition | Gene-family content tree | Sequence-based tree | Whole-genome tree reported by Fontaine et al. (2015) | X-chromosome topology reported by Fontaine et al. (2015) | Species tree reported by Thawornwattana et al. (2018) | Published network reported by Fontaine et al. (2015) | Published network reported by Thawornwattana et al. (2018) |
|---|---|---|---|---|---|---|---|---|
| odd | LNK1 | -704142.34 | -704202.35 | -704338.24 | -704678.21 | -704634.17 | -705051.38 | -704733.94 |
| | LNK2 | -686875.47 | -687017.15 | -687106.5 | -687388.98 | -687347.1 | -687626.93 | -687604.17 |
| even | LNK3 | -701702.11 | -701739.77 | -701826.31 | -702121.56 | -702105.55 | -702441.95 | -702321.81 |
| | LNK4 | -660138.73 | -660205.33 | -660252.5 | -660558.52 | -660427.67 | -660755.97 | -660753.06 |

### Table S12. Posterior parameter estimates for the best-supported network inferred under the gene-family content species tree.

**A. Posterior parameter estimates**

| node | τ | Time (Ma) | 95% HPD of τ | θ | 95% HPD of θ |
|---|---|---|---|---|---|
| N6 | 0.020982 | 0.6812 | 0.020465-0.021514 | 0.016012 | 0.015036-0.016987 |
| N5 | 0.018135 | 0.5888 | 0.017325-0.018959 | 0.003799 | 0.002774-0.004774 |
| N4 | 0.011888 | 0.3860 | 0.011667-0.012111 | 0.011502 | 0.010432-0.012580 |
| Z1 | 0.011874 | 0.3855 | 0.011650-0.012098 | 0.189744 | 0.147390-0.234417 |
| N3 | 0.011874 | 0.3855 | 0.011650-0.012098 | 0.013082 | 0.001327-0.028420 |
| Z5 | 0.011269 | 0.3659 | 0.010257-0.012022 | 0.009604 | 0.000167-0.023353 |
| Z4 | 0.009656 | 0.3135 | 0.009381-0.009928 | 0.007692 | 0.001565-0.015602 |
| Z3 | 0.009656 | 0.3135 | 0.009381-0.009928 | 0.014052 | 0.012252-0.015880 |
| Z9 | 0.009541 | 0.3098 | 0.009187-0.009876 | 0.010491 | 0.001098-0.022521 |
| N2 | 0.006717 | 0.2181 | 0.006479-0.006957 | 0.012616 | 0.011338-0.013900 |
| Z11 | 0.003592 | 0.1166 | 0.003138-0.003979 | 0.004860 | 0.004207-0.005540 |
| Z12 | 0.003592 | 0.1166 | 0.003138-0.003979 | 0.014439 | 0.013138-0.015738 |
| N1 | 0.002681 | 0.0870 | 0.002461-0.002930 | 0.018461 | 0.011084-0.025018 |
| Z7 | 0.002670 | 0.0867 | 0.002444-0.002918 | 0.012807 | 0.001165-0.027856 |
| Z8 | 0.002670 | 0.0867 | 0.002444-0.002918 | 0.004957 | 0.004401-0.005507 |
| Z10 | 0.002302 | 0.0747 | 0.001900-0.002655 | 0.021385 | 0.008041-0.037616 |

**B. Savage–Dickey density-ratio tests for hybrid speciation**

| hybrid speciation | Bε (ε=10-5) | Bε (ε=10-6) |
|---|---|---|
| TauN1-TauZ8 | 0.004 | 0.003 |

The corresponding positions of all node labels in the MSci network are shown in Fig. S3B

### Table S13. D-statistic results for data simulated under the gene-family content species-tree topology

| P1 | P2 | P3 | D | Z-score | p-value | f4-ratio | BBAA | ABBA | BABA | Dp | adjusted_p_value |
|---|---|---|---|---|---|---|---|---|---|---|---|
| B | C | F | 0.564587 | 55.9618 | 2.3e-16 | 0.328121 | 2818.25 | 8089.5 | 2251.25 | 0.44367 | 4.6e-15 |
| B | D | F | 0.565029 | 51.0062 | 2.3e-16 | 0.326814 | 2829.75 | 8053.25 | 2238.25 | 0.443174 | 4.6e-15 |
| B | E | F | 0.546915 | 45.2949 | 2.3e-16 | 0.313255 | 2807.5 | 7882.5 | 2308.75 | 0.428791 | 4.6e-15 |
| A | C | F | 0.523067 | 47.5981 | 2.3e-16 | 0.289896 | 2966.12 | 7097.88 | 2222.62 | 0.396794 | 4.6e-15 |
| A | D | F | 0.524526 | 41.0935 | 2.3e-16 | 0.288513 | 2990.88 | 7051.12 | 2199.12 | 0.396369 | 4.6e-15 |
| A | E | F | 0.5017 | 41.9078 | 2.3e-16 | 0.274168 | 2983.75 | 6900.5 | 2289.75 | 0.378737 | 4.6e-15 |
| B | A | D | 0.108878 | 6.86181 | 6.79913e-12 | 0.0699573 | 2850.12 | 5923.62 | 4760.38 | 0.0859487 | 1.35983e-10 |
| B | A | E | 0.108858 | 6.77766 | 1.2214e-11 | 0.0647537 | 2923.38 | 5899.12 | 4740.88 | 0.0853946 | 2.4428e-10 |
| B | A | C | 0.107023 | 6.3318 | 2.4232e-10 | 0.0692137 | 2868.12 | 5893.38 | 4753.88 | 0.0843114 | 4.8464e-09 |
| B | A | F | 0.0935587 | 5.4869 | 4.09057e-08 | 0.0541224 | 3346.25 | 5628 | 4665 | 0.0706051 | 8.18114e-07 |
| D | C | E | 0.0984036 | 4.55074 | 5.34582e-06 | 0.0464084 | 4292.75 | 2305 | 1892 | 0.0486469 | 0.000106916 |
| E | C | F | 0.0660589 | 6.19404 | 5.86421e-10 | 0.0216794 | 6202 | 2134.25 | 1869.75 | 0.0259161 | 1.17284e-08 |
| E | D | F | 0.0579023 | 3.75819 | 0.000171149 | 0.0197738 | 5881.88 | 2203.88 | 1962.62 | 0.0240098 | 0.00342298 |
| Taxa were labeled A–F, corresponding respectively to A. melas, A. merus, A. gambiae, A. coluzzii, A. arabiensis, and A. quadriannulatus | | | | | | | | | | | |

### Table S14. Posterior parameter estimates for the simulated dataset

| A. Posterior parameter estimates | | | | | |
|---|---|---|---|---|---|
| **node** | **τ** | **Time (Ma)** | **95% HPD of τ** | **θ** | **95% HPD of θ** |
| N6 | 0.020515 | 0.6661 | 0.019966-0.021090 | 0.016328 | 0.015468-0.017181 |
| N5 | 0.018455 | 0.5992 | 0.017700-0.019218 | 0.003006 | 0.001668-0.004298 |
| N4 | 0.011920 | 0.3870 | 0.011745-0.012091 | 0.010931 | 0.010085-0.011810 |
| Z1 | 0.011866 | 0.3853 | 0.011653-0.012065 | 0.080966 | 0.059102-0.104720 |
| N3 | 0.011866 | 0.3853 | 0.011653-0.012065 | 0.014336 | 0.001428-0.030455 |
| Z5 | 0.010191 | 0.3309 | 0.008904-0.011608 | 0.009439 | 0.000154-0.023037 |
| Z4 | 0.009589 | 0.3113 | 0.009312-0.009869 | 0.009565 | 0.000177-0.023041 |
| Z3 | 0.009589 | 0.3113 | 0.009312-0.009869 | 0.014293 | 0.011546-0.017004 |
| Z9 | 0.008979 | 0.2915 | 0.008245-0.009628 | 0.019156 | 0.003824-0.035070 |
| N2 | 0.006848 | 0.2223 | 0.006577-0.007111 | 0.011203 | 0.008018-0.013942 |
| Z11 | 0.003473 | 0.1128 | 0.003269-0.003677 | 0.004982 | 0.004495-0.005461 |
| Z12 | 0.003473 | 0.1128 | 0.003269-0.003677 | 0.015682 | 0.014120-0.017269 |
| N1 | 0.003001 | 0.0974 | 0.002781-0.003206 | 0.008767 | 0.003444-0.014340 |
| Z7 | 0.002969 | 0.0964 | 0.002749-0.003175 | 0.014940 | 0.001513-0.031756 |
| Z8 | 0.002969 | 0.0964 | 0.002749-0.003175 | 0.004353 | 0.003831-0.004879 |
| Z10 | 0.002179 | 0.0707 | 0.001159-0.003031 | 0.012588 | 0.000678-0.027710 |
| **B. Introgression probabilities** | | | | | |
| **Gene-flow event** | | **B10 (ε=0.01)** | | **φ** | |
| Z1→Z2 | | ∞ | | 0.41233 | |
| Z3→Z4 | | ∞ | | 0.811 | |
| Z6→Z5 | | ∞ | | 0.3214 | |
| Z7→Z8 | | ∞ | | 0.1396 | |
| Z8→Z7 | | ∞ | | 0.0412 | |
| Z9→Z10 | | ∞ | | 0.0904 | |
| Z12→Z11 | | ∞ | | 0.0997 | |

### Table S15. Posterior parameter estimates for the reduced three-species MSci network

| A. Posterior parameter estimates | | | | | |
|---|---|---|---|---|---|
| **node** | **τ** | **Time (Ma)** | **95% HPD of τ** | **θ** | **95% HPD of θ** |
| M | 0.016289 | 0.5289 | 0.014859-0.017820 | 0.016048 | 0.013690-0.018535 |
| T | 0.007793 | 0.2530 | 0.007554-0.008029 | 0.007609 | 0.007064-0.008138 |
| X | 0.002814 | 0.0914 | 0.002689-0.002938 | 0.014748 | 0.013754-0.015731 |
| P | 0.002808 | 0.0912 | 0.002679-0.002930 | 0.001348 | 0.000198-0.003402 |
| Q | 0.002808 | 0.0912 | 0.002679-0.002930 | 0.004927 | 0.004398-0.005454 |
| J | 0.002805 | 0.0911 | 0.002677-0.002928 | 0.001219 | 0.000196-0.003044 |
| K | 0.002805 | 0.0911 | 0.002677-0.002928 | 0.001230 | 0.000194-0.003090 |
| N | 0.002633 | 0.0855 | 0.002060-0.002930 | 0.066942 | 0.000173-0.169031 |
| **B. Introgression probabilities** | | | | | |
| **Gene-flow event** | | **B10 (ε=0.01)** | | **φ** | |
| M→N | | ∞ | | 0.02 | |
| Q→P | | ∞ | | 0.05 | |
| J→K | | ∞ | | 0.15 | |
| **C. Savage–Dickey density-ratio tests for hybrid speciation** | | | | | |
| **hybrid speciation** | | **Bε (ε=10-5)** | | **Bε (ε=10-6)** | |
| TauX-TauP | | 0.0102 | | 0.006 | |

**Table S16. Savage–Dickey density-ratio tests for 10 simulated datasets under the reduced three-species model**

| Replicate | Bε (ε = $10^{-5}$) | Bε (ε = $10^{-6}$) |
|---|---|---|
| 1 | 0.0120456507145 | 0.003507565337 |
| 2 | 0.0103066111526 | 0.00599726857075 |
| 3 | 0.0130047635608 | 0.0234234234234 |
| 4 | 0.01166821458 | 0.0124640460211 |
| 5 | 0.0252310717797 | 0.0228658536585 |
| 6 | 0.0105878538572 | 0.00845779493204 |
| 7 | 0.0230254350736 | 0.0239407651172 |
| 8 | 0.00761389969021 | 0.00224519632414 |
| 9 | 0.00565964681918 | 0.00514693674249 |
| 10 | 0.00799653535025 | 0.00273534269199 |

### Table S17. HKA test results for comparison-specific XP-CLR candidate intervals in *Anopheles gambiae*

| Interval ID | Candidate ratio | Neutral-background ratio | HKA class | P value | Comparison | Adjusted P value | Chr. | Start (bp) | End (bp) |
|---|---|---|---|---|---|---|---|---|---|
| colaa | 53.42857143 | 94.97383178 | NotSig | 0.125793349 | ara2col | 1 | 2L | 4100001 | 4120000 |
| colab | 56.875 | 94.97383178 | PosSelect | 0.007720081 | ara2col | 0.208442174 | 2L | 5680001 | 5760000 |
| colac | 143.4561404 | 94.97383178 | NegSelect | 0.002240384 | ara2col | 0.060490371 | 2L | 9380001 | 9800000 |
| colad | Inf | 94.97383178 | NegSelect | 0.00520526 | ara2col | 0.14054201 | 2R | 8300001 | 8320000 |
| colae | Inf | 94.97383178 | NegSelect | 0.011485694 | ara2col | 0.310113744 | 2R | 38400001 | 38420000 |
| colaf | 652.5172414 | 94.97383178 | NegSelect | 1.37E-41 | ara2col | 3.71E-40 | 2R | 48820001 | 49440000 |
| colag | 343.3157895 | 94.97383178 | NegSelect | 2.18E-11 | ara2col | 5.88E-10 | 3R | 28420001 | 28660000 |
| colah | 148.2068966 | 94.97383178 | NegSelect | 0.017456751 | ara2col | 0.471332286 | 3R | 41280001 | 41400000 |
| colai | 55.94890511 | 94.97383178 | PosSelect | 2.06E-07 | ara2col | 5.57E-06 | 3R | 48860001 | 49480000 |
| colaj | 8.831111111 | 94.97383178 | PosSelect | 7.17E-127 | ara2col | 1.94E-125 | 3R | 52040001 | 52620000 |
| araaa | 6.742424242 | 43.35518906 | PosSelect | 2.23E-29 | col2ara | 6.02E-28 | 2L | 2420001 | 2580000 |
| araab | 58.69230769 | 43.35518906 | NotSig | 0.330710426 | col2ara | 1 | 2L | 3920001 | 3940000 |
| araac | 159.1666667 | 43.35518906 | NegSelect | 0.000161268 | col2ara | 0.004354244 | 2L | 5660001 | 5680000 |
| araad | 295.4090909 | 43.35518906 | NegSelect | 7.56E-70 | col2ara | 2.04E-68 | 2L | 12980001 | 13300000 |
| araae | 303.3333333 | 43.35518906 | NegSelect | 2.90E-06 | col2ara | 7.84E-05 | 2R | 37740001 | 37760000 |
| araaf | 26.84615385 | 43.35518906 | PosSelect | 0.022948416 | col2ara | 0.619607233 | 2R | 42740001 | 42760000 |
| araag | Inf | 43.35518906 | NegSelect | 7.50E-06 | col2ara | 0.000202519 | 2R | 50460001 | 50480000 |
| araah | 195.5 | 43.35518906 | NegSelect | 0.000227017 | col2ara | 0.006129464 | 2R | 56480001 | 56500000 |
| araai | 62.98557692 | 43.35518906 | NegSelect | 1.24E-17 | col2ara | 3.35E-16 | 2R | 57640001 | 60340000 |
| araaj | 23.70409712 | 43.35518906 | PosSelect | 2.14E-37 | col2ara | 5.79E-36 | 3L | 840001 | 2520000 |
| araak | 19.95078031 | 43.35518906 | PosSelect | 7.41E-71 | col2ara | 2.00E-69 | 3L | 4820001 | 5740000 |
| araal | 100.9662921 | 43.35518906 | NegSelect | 8.20E-19 | col2ara | 2.21E-17 | 3L | 7060001 | 7840000 |
| araam | 23.07692308 | 43.35518906 | PosSelect | 0.033641139 | col2ara | 0.908310752 | 3L | 41880001 | 41900000 |
| araan | 164.5652174 | 43.35518906 | NegSelect | 2.42E-15 | col2ara | 6.54E-14 | 3R | 37760001 | 37840000 |
| araao | 83.69230769 | 43.35518906 | NegSelect | 0.013669902 | col2ara | 0.369087348 | 3R | 42280001 | 42300000 |
| araap | 40.68852459 | 43.35518906 | NotSig | 0.636095719 | col2ara | 1 | 3R | 48380001 | 48440000 |
| araaq | 166.4444444 | 43.35518906 | NegSelect | 6.58E-07 | col2ara | 1.78E-05 | 3R | 50100001 | 50140000 |

Comparison labels denote the putative ancestry contrast used in the XP-CLR analysis. PosSelect, NegSelect, and NotSig denote positive-selection, negative-selection, and nonsignificant HKA classifications, respectively.

### Table S18. Genes of HKA-supported XP-CLR candidate intervals consistent with *A. coluzzii* or *A. arabiensis* ancestry in *A. gambiae*

| Sequence | Source | Feature | Start | End | Strand | Attributes |
|---|---|---|---|---|---|---|
| AgamP4_2L | VEuPathDB | protein_coding_gene | 2358158 | 2431617 | + | ID=AGAP004707;Name=para;description=voltage-gated sodium channel;ebi_biotype=protein_coding |
| AgamP4_2L | VEuPathDB | protein_coding_gene | 2471589 | 2474401 | - | ID=AGAP004708;description=arginyl-tRNA synthetase;ebi_biotype=protein_coding |
| AgamP4_2L | VEuPathDB | protein_coding_gene | 2482553 | 2483310 | + | ID=AGAP004709;Name=mRpL18;description=39S ribosomal protein L18, mitochondrial;ebi_biotype=protein_coding |
| AgamP4_2L | VEuPathDB | protein_coding_gene | 2483226 | 2483631 | - | ID=AGAP004710;description=ubiquinol-cytochrome c reductase subunit 9;ebi_biotype=protein_coding |
| AgamP4_2L | VEuPathDB | protein_coding_gene | 2487698 | 2489698 | + | ID=AGAP004711;description=ATP-dependent RNA helicase DDX41;ebi_biotype=protein_coding |
| AgamP4_2L | VEuPathDB | protein_coding_gene | 2506572 | 2507341 | + | ID=AGAP004712;description=unspecified product;ebi_biotype=protein_coding |
| AgamP4_2L | VEuPathDB | protein_coding_gene | 2558997 | 2560250 | - | ID=AGAP004713;description=PHD-type domain-containing protein [Source:UniProtKB/TrEMBL;Acc:A0A1S4GLV7];ebi_biotype=protein_coding |
| AgamP4_2L | VEuPathDB | protein_coding_gene | 2567155 | 2574572 | + | ID=AGAP004714;description=unspecified product;ebi_biotype=protein_coding |
| AgamP4_3R | VEuPathDB | protein_coding_gene | 48812145 | 48885340 | - | ID=AGAP010090;description=receptor-type tyrosine-protein phosphatase F;ebi_biotype=protein_coding |
| AgamP4_3R | VEuPathDB | protein_coding_gene | 49065386 | 49069554 | + | ID=AGAP010094;description=nonmuscle myosin heavy chain-A;ebi_biotype=protein_coding |
| AgamP4_3R | VEuPathDB | protein_coding_gene | 49071967 | 49073130 | + | ID=AGAP010095;Name=CPR82;description=cuticular protein RR-2 family 82;ebi_biotype=protein_coding |
| AgamP4_3R | VEuPathDB | protein_coding_gene | 49106142 | 49113838 | - | ID=AGAP010096;description=unspecified product;ebi_biotype=protein_coding |
| AgamP4_3R | VEuPathDB | protein_coding_gene | 49126555 | 49127312 | - | ID=AGAP010097;Name=CPR107;description=cuticular protein RR-2 family 107;ebi_biotype=protein_coding |
| AgamP4_3R | VEuPathDB | protein_coding_gene | 49131810 | 49132540 | - | ID=AGAP010098;Name=CPR83;description=cuticular protein RR-2 family 83;ebi_biotype=protein_coding |
| AgamP4_3R | VEuPathDB | protein_coding_gene | 49136221 | 49136690 | - | ID=AGAP010099;Name=CPR108;description=cuticular protein RR-2 family 108;ebi_biotype=protein_coding |
| AgamP4_3R | VEuPathDB | protein_coding_gene | 49137589 | 49138196 | + | ID=AGAP010100;Name=CPR84;description=cuticular protein RR-2 family 84;ebi_biotype=protein_coding |
| AgamP4_3R | VEuPathDB | protein_coding_gene | 49142195 | 49142795 | - | ID=AGAP010101;Name=CPR85;description=cuticular protein RR-2 family 85;ebi_biotype=protein_coding |
| AgamP4_3R | VEuPathDB | protein_coding_gene | 49150129 | 49150590 | + | ID=AGAP010103;Name=CPR86;description=cuticular protein RR-2 family 86;ebi_biotype=protein_coding |
| AgamP4_3R | VEuPathDB | protein_coding_gene | 49155284 | 49155745 | + | ID=AGAP010104;Name=CPR87;description=cuticular protein RR-2 family 87;ebi_biotype=protein_coding |
| AgamP4_3R | VEuPathDB | protein_coding_gene | 49157894 | 49158335 | + | ID=AGAP010105;Name=CPR88;description=cuticular protein RR-2 family 88;ebi_biotype=protein_coding |
| AgamP4_3R | VEuPathDB | protein_coding_gene | 49160573 | 49161034 | + | ID=AGAP010106;Name=CPR89;description=cuticular protein RR-2 family 89;ebi_biotype=protein_coding |
| AgamP4_3R | VEuPathDB | protein_coding_gene | 49164519 | 49164980 | + | ID=AGAP010107;Name=CPR90;description=cuticular protein RR-2 family 90;ebi_biotype=protein_coding |
| AgamP4_3R | VEuPathDB | protein_coding_gene | 49169540 | 49169977 | + | ID=AGAP010108;Name=CPR91;description=cuticular protein RR-2 family 91;ebi_biotype=protein_coding |
| AgamP4_3R | VEuPathDB | protein_coding_gene | 49182656 | 49183063 | + | ID=AGAP010109;Name=CPR150;description=cuticular protein 150;ebi_biotype=protein_coding |
| AgamP4_3R | VEuPathDB | protein_coding_gene | 49185444 | 49187681 | + | ID=AGAP010110;description=N-acetyltransferase domain-containing protein [Source:UniProtKB/TrEMBL;Acc:A0A1S4H2Y8];ebi_biotype=protein_coding |
| AgamP4_3R | VEuPathDB | protein_coding_gene | 49188351 | 49190046 | + | ID=AGAP010111;description=N-acetyltransferase domain-containing protein [Source:UniProtKB/TrEMBL;Acc:A0A1S4H305];ebi_biotype=protein_coding |
| AgamP4_3R | VEuPathDB | protein_coding_gene | 49190235 | 49191000 | - | ID=AGAP010112;Name=CPR92;description=cuticular protein RR-2 family 92;ebi_biotype=protein_coding |
| AgamP4_3R | VEuPathDB | protein_coding_gene | 49193017 | 49193877 | - | ID=AGAP010113;Name=CPR93;description=cuticular protein RR-2 family 93;ebi_biotype=protein_coding |
| AgamP4_3R | VEuPathDB | protein_coding_gene | 49196276 | 49197137 | - | ID=AGAP010114;Name=CPR94;description=cuticular protein RR-2 family 94;ebi_biotype=protein_coding |
| AgamP4_3R | VEuPathDB | protein_coding_gene | 49201849 | 49207894 | + | ID=AGAP010115;description=unspecified product;ebi_biotype=protein_coding |
| AgamP4_3R | VEuPathDB | protein_coding_gene | 49210876 | 49211678 | + | ID=AGAP010116;Name=CPR109;description=cuticular protein RR-2 family 109;ebi_biotype=protein_coding |
| AgamP4_3R | VEuPathDB | protein_coding_gene | 49215715 | 49216528 | + | ID=AGAP010117;Name=CPR95;description=cuticular protein RR-2 family 95;ebi_biotype=protein_coding |
| AgamP4_3R | VEuPathDB | protein_coding_gene | 49216717 | 49217949 | - | ID=AGAP010118;description=N-acetyltransferase domain-containing protein [Source:UniProtKB/TrEMBL;Acc:A0A1S4H3W9];ebi_biotype=protein_coding |
| AgamP4_3R | VEuPathDB | protein_coding_gene | 49220511 | 49221282 | + | ID=AGAP010119;Name=CPR96;description=cuticular protein RR-2 family 96;ebi_biotype=protein_coding |
| AgamP4_3R | VEuPathDB | protein_coding_gene | 49227988 | 49228876 | + | ID=AGAP010120;Name=CPR97;description=cuticular protein RR-2 family 97;ebi_biotype=protein_coding |

| Sequence | Source | Feature | Start | End | Strand | Attributes |
|---|---|---|---|---|---|---|
| AgamP4_3R | VEuPathDB | protein_coding_gene | 49230063 | 49230500 | + | ID=AGAP010121;Name=CPR149;description=cuticular protein 149;ebi_biotype=protein_coding |
| AgamP4_3R | VEuPathDB | protein_coding_gene | 49237379 | 49238739 | + | ID=AGAP010122;Name=CPR132;description=cuticular protein RR-2 family 132;ebi_biotype=protein_coding |
| AgamP4_3R | VEuPathDB | protein_coding_gene | 49239817 | 49240669 | + | ID=AGAP010123;Name=CPR131;description=cuticular protein RR-2 family 131;ebi_biotype=protein_coding |
| AgamP4_3R | VEuPathDB | protein_coding_gene | 49242946 | 49243750 | + | ID=AGAP010124;Name=CPR98;description=cuticular protein RR-2 family 98;ebi_biotype=protein_coding |
| AgamP4_3R | VEuPathDB | protein_coding_gene | 49249463 | 49251287 | + | ID=AGAP010125;description=N-acetyltransferase domain-containing protein [Source:UniProtKB/TrEMBL;Acc:A0A1S4H303];ebi_biotype=protein_coding |
| AgamP4_3R | VEuPathDB | protein_coding_gene | 49251731 | 49252603 | - | ID=AGAP010126;Name=CPR142;description=cuticular protein RR-2 family 142;ebi_biotype=protein_coding |
| AgamP4_3R | VEuPathDB | protein_coding_gene | 49254604 | 49255464 | - | ID=AGAP010127;Name=CPR99;description=cuticular protein RR-2 family 99;ebi_biotype=protein_coding |
| AgamP4_3R | VEuPathDB | protein_coding_gene | 49260060 | 49260851 | + | ID=AGAP010128;Name=CPR100;description=cuticular protein RR-2 family 100;ebi_biotype=protein_coding |
| AgamP4_3R | VEuPathDB | protein_coding_gene | 49261530 | 49263906 | - | ID=AGAP010129;description=unspecified product;ebi_biotype=protein_coding |
| AgamP4_3R | VEuPathDB | protein_coding_gene | 49265590 | 49267033 | - | ID=AGAP010130;description=3,2-trans-enoyl-CoA isomerase mitochondrial;ebi_biotype=protein_coding |
| AgamP4_3R | VEuPathDB | protein_coding_gene | 49268823 | 49275992 | + | ID=AGAP010131;description=ornithine decarboxylase antizyme 1;ebi_biotype=protein_coding |
| AgamP4_3R | VEuPathDB | protein_coding_gene | 49285621 | 49287337 | + | ID=AGAP010132;Name=SCRBQ1;description=Class B Scavenger Receptor (CD36 domain).;ebi_biotype=protein_coding |
| AgamP4_3R | VEuPathDB | protein_coding_gene | 49291876 | 49295766 | + | ID=AGAP010133;Name=SCRBQ2;description=Class B Scavenger Receptor (CD36 domain).;ebi_biotype=protein_coding |
| AgamP4_3R | VEuPathDB | protein_coding_gene | 49302810 | 49305364 | + | ID=AGAP010134;description=arrestin-1;ebi_biotype=protein_coding |
| AgamP4_3R | VEuPathDB | protein_coding_gene | 49308809 | 49314974 | - | ID=AGAP010135;description=NCK adaptor protein;ebi_biotype=protein_coding |
| AgamP4_3R | VEuPathDB | protein_coding_gene | 49317276 | 49317725 | - | ID=AGAP010136;description=PfkB domain-containing protein [Source:UniProtKB/TrEMBL;Acc:A0A1S4H3Z0];ebi_biotype=protein_coding |
| AgamP4_3R | VEuPathDB | protein_coding_gene | 49319449 | 49321593 | - | ID=AGAP010137;description=adenosine kinase;ebi_biotype=protein_coding |
| AgamP4_3R | VEuPathDB | protein_coding_gene | 49324570 | 49350676 | + | ID=AGAP010138;description=uncharacterized protein yjbQ;ebi_biotype=protein_coding |
| AgamP4_3R | VEuPathDB | protein_coding_gene | 49356301 | 49362525 | - | ID=AGAP010139;description=GMP synthase (glutamine-hydrolysing);ebi_biotype=protein_coding |
| AgamP4_3R | VEuPathDB | protein_coding_gene | 49362915 | 49364048 | - | ID=AGAP010140;description=hydrolases of HD superfamily;ebi_biotype=protein_coding |
| AgamP4_3R | VEuPathDB | protein_coding_gene | 49364451 | 49365588 | + | ID=AGAP010141;description=DnaJ homolog subfamily C member 4;ebi_biotype=protein_coding |
| AgamP4_3R | VEuPathDB | protein_coding_gene | 49370055 | 49390808 | - | ID=AGAP010142;Name=Dat;description=dopamine N-acetyltransferase;ebi_biotype=protein_coding |
| AgamP4_3R | VEuPathDB | protein_coding_gene | 49409660 | 49414135 | + | ID=AGAP010143;description=BED-type domain-containing protein [Source:UniProtKB/TrEMBL;Acc:A0A1S4H324];ebi_biotype=protein_coding |
| AgamP4_3R | VEuPathDB | protein_coding_gene | 49420819 | 49421462 | + | ID=AGAP010144;description=unspecified product;ebi_biotype=protein_coding |
| AgamP4_3R | VEuPathDB | protein_coding_gene | 49425840 | 49433965 | - | ID=AGAP010145;description=Yellow [Source:UniProtKB/TrEMBL;Acc:A0A1S4H3W2];ebi_biotype=protein_coding |
| AgamP4_3R | VEuPathDB | protein_coding_gene | 49434762 | 49436679 | - | ID=AGAP010146;description=unspecified product;ebi_biotype=protein_coding |
| AgamP4_3R | VEuPathDB | protein_coding_gene | 49458955 | 49484075 | + | ID=AGAP010147;description=myosin heavy chain;ebi_biotype=protein_coding |
| AgamP4_3R | VEuPathDB | protein_coding_gene | 52042711 | 52059885 | + | ID=AGAP010283;description=wingless-type MMTV integration site family, member 7;ebi_biotype=protein_coding |
| AgamP4_3R | VEuPathDB | protein_coding_gene | 52067738 | 52115209 | + | ID=AGAP010286;description=unspecified product;ebi_biotype=protein_coding |
| AgamP4_3R | VEuPathDB | protein_coding_gene | 52161878 | 52162651 | + | ID=AGAP010287;description=unspecified product;ebi_biotype=protein_coding |
| AgamP4_3R | VEuPathDB | protein_coding_gene | 52172251 | 52173633 | + | ID=AGAP010288;description=YY1-associated factor 2;ebi_biotype=protein_coding |
| AgamP4_3R | VEuPathDB | protein_coding_gene | 52210677 | 52214536 | - | ID=AGAP010289;description=F-box and leucine-rich repeat protein 14 [Source:UniProtKB/TrEMBL;Acc:A0A1S4H4H8];ebi_biotype=protein_coding |
| AgamP4_3R | VEuPathDB | protein_coding_gene | 52228421 | 52248901 | + | ID=AGAP010290;description=wingless-type MMTV integration site family, member 5;ebi_biotype=protein_coding |
| AgamP4_3R | VEuPathDB | protein_coding_gene | 52265896 | 52271136 | - | ID=AGAP010291;description=unspecified product;ebi_biotype=protein_coding |
| AgamP4_3R | VEuPathDB | protein_coding_gene | 52301033 | 52349893 | - | ID=AGAP010292;description=guanine nucleotide exchange factor VAV;ebi_biotype=protein_coding |
| AgamP4_3R | VEuPathDB | protein_coding_gene | 52352368 | 52361114 | + | ID=AGAP010293;Name=GC1;description=mitochondrial glutamate carrier 1;ebi_biotype=protein_coding |
| AgamP4_3R | VEuPathD | protein_coding_g | 52418651 | 52422892 | - | ID=AGAP010294;description=unspecified product;ebi_biotype=protein_coding |

| Sequence | Source | Feature | Start | End | Strand | Attributes |
|---|---|---|---|---|---|---|
| | B | ene | | | | |
| AgamP4_3R | VEuPathDB | protein_coding_gene | 52456487 | 52554377 | - | ID=AGAP010295;description=Ca_chan_IQ domain-containing protein [Source:UniProtKB/TrEMBL;Acc:A0A1S4H3K3];ebi_biotype=protein_coding |
| AgamP4_3R | VEuPathDB | protein_coding_gene | 52614350 | 52617318 | + | ID=AGAP010297;description=unspecified product;ebi_biotype=protein_coding |
| AgamP4_3L | VEuPathDB | protein_coding_gene | 850534 | 859465 | - | ID=AGAP010326;Name=AQP3;description=aquaporin;ebi_biotype=protein_coding |
| AgamP4_3L | VEuPathDB | protein_coding_gene | 869455 | 875290 | - | ID=AGAP010327;description=SH3-binding domain kinase;ebi_biotype=protein_coding |
| AgamP4_3L | VEuPathDB | protein_coding_gene | 884186 | 885693 | + | ID=AGAP010328;description=unspecified product;ebi_biotype=protein_coding |
| AgamP4_3L | VEuPathDB | protein_coding_gene | 984808 | 985451 | + | ID=AGAP010329;description=unspecified product;ebi_biotype=protein_coding |
| AgamP4_3L | VEuPathDB | protein_coding_gene | 1016366 | 1017211 | - | ID=AGAP010330;description=unspecified product;ebi_biotype=protein_coding |
| AgamP4_3L | VEuPathDB | protein_coding_gene | 1237241 | 1252839 | + | ID=AGAP010331;description=heat shock protein 110kDa;ebi_biotype=protein_coding |
| AgamP4_3L | VEuPathDB | protein_coding_gene | 1262900 | 1265820 | + | ID=AGAP010332;description=C2H2-type domain-containing protein [Source:UniProtKB/TrEMBL;Acc:A0A1S4H4H6];ebi_biotype=protein_coding |
| AgamP4_3L | VEuPathDB | protein_coding_gene | 1619302 | 1621822 | + | ID=AGAP010334;description=glucosyl/glucuronosyl transferases;ebi_biotype=protein_coding |
| AgamP4_3L | VEuPathDB | protein_coding_gene | 1566392 | 1659715 | + | ID=AGAP010335;description=NEL-LIKE PROTEIN 1;ebi_biotype=protein_coding |
| AgamP4_3L | VEuPathDB | protein_coding_gene | 1661447 | 1663508 | - | ID=AGAP010336;description=tubulin polyglutamylase TTLL1;ebi_biotype=protein_coding |
| AgamP4_3L | VEuPathDB | protein_coding_gene | 1664219 | 1665110 | - | ID=AGAP010337;description=ubiquinol-cytochrome c reductase subunit 8;ebi_biotype=protein_coding |
| AgamP4_3L | VEuPathDB | protein_coding_gene | 1687955 | 1688766 | + | ID=AGAP010338;Name=mRpL23;description=39S ribosomal protein L23, mitochondrial;ebi_biotype=protein_coding |
| AgamP4_3L | VEuPathDB | protein_coding_gene | 1689490 | 1690978 | - | ID=AGAP010339;description=WW domain-binding protein 4;ebi_biotype=protein_coding |
| AgamP4_3L | VEuPathDB | protein_coding_gene | 1727868 | 1787423 | - | ID=AGAP010340;description=unspecified product;ebi_biotype=protein_coding |
| AgamP4_3L | VEuPathDB | protein_coding_gene | 1792937 | 1794099 | - | ID=AGAP010341;description=testis-specific serine kinase;ebi_biotype=protein_coding |
| AgamP4_3L | VEuPathDB | protein_coding_gene | 1804512 | 1866030 | - | ID=AGAP010342;description=unspecified product;ebi_biotype=protein_coding |
| AgamP4_3L | VEuPathDB | protein_coding_gene | 1877545 | 1889368 | + | ID=AGAP010343;description=CTL-like protein 2;ebi_biotype=protein_coding |
| AgamP4_3L | VEuPathDB | protein_coding_gene | 1894411 | 1899646 | + | ID=AGAP010344;description=solute carrier family 26, other;ebi_biotype=protein_coding |
| AgamP4_3L | VEuPathDB | protein_coding_gene | 1903438 | 1912215 | - | ID=AGAP010345;description=OTU domain-containing protein 7;ebi_biotype=protein_coding |
| AgamP4_3L | VEuPathDB | protein_coding_gene | 1913738 | 1918285 | + | ID=AGAP010346;Name=Mer;description=merlin (moesin-ezrin-radixin-like protein);ebi_biotype=protein_coding |
| AgamP4_3L | VEuPathDB | protein_coding_gene | 1918545 | 1920511 | + | ID=AGAP010347;Name=CuSOD3;description=copper-zinc superoxide dismutase 3;ebi_biotype=protein_coding |
| AgamP4_3L | VEuPathDB | protein_coding_gene | 1920561 | 1922286 | - | ID=AGAP010348;description=unspecified product;ebi_biotype=protein_coding |
| AgamP4_3L | VEuPathDB | protein_coding_gene | 1937808 | 1941764 | - | ID=AGAP010349;description=unspecified product;ebi_biotype=protein_coding |
| AgamP4_3L | VEuPathDB | protein_coding_gene | 1948270 | 1950111 | + | ID=AGAP010350;description=unspecified product;ebi_biotype=protein_coding |
| AgamP4_3L | VEuPathDB | protein_coding_gene | 1950125 | 1954253 | - | ID=AGAP010351;description=insulysin;ebi_biotype=protein_coding |
| AgamP4_3L | VEuPathDB | protein_coding_gene | 1954505 | 1955821 | + | ID=AGAP010352;description=DNA/RNA-binding protein KIN17;ebi_biotype=protein_coding |
| AgamP4_3L | VEuPathDB | protein_coding_gene | 1959754 | 1962699 | + | ID=AGAP010353;description=DNA cross-link repair 1A protein;ebi_biotype=protein_coding |
| AgamP4_3L | VEuPathDB | protein_coding_gene | 2059637 | 2061051 | - | ID=AGAP010354;description=osiris 21;ebi_biotype=protein_coding |
| AgamP4_3L | VEuPathDB | protein_coding_gene | 2082866 | 2113040 | + | ID=AGAP010355;description=unspecified product;ebi_biotype=protein_coding |
| AgamP4_3L | VEuPathDB | protein_coding_gene | 2168978 | 2175293 | - | ID=AGAP010358;description=paired box protein 3/7;ebi_biotype=protein_coding |
| AgamP4_3L | VEuPathDB | protein_coding_gene | 2202981 | 2221987 | + | ID=AGAP010359;description=Pairberry;ebi_biotype=protein_coding |
| AgamP4_3L | VEuPathDB | protein_coding_gene | 2238417 | 2239634 | - | ID=AGAP010362;description=unspecified product;ebi_biotype=protein_coding |
| AgamP4_3L | VEuPathDB | protein_coding_gene | 2242724 | 2243715 | + | ID=AGAP010363;description=unspecified product;ebi_biotype=protein_coding |
| AgamP4_3L | VEuPathDB | protein_coding_gene | 2246781 | 2247865 | + | ID=AGAP010364;description=unspecified product;ebi_biotype=protein_coding |

| Sequence | Source | Feature | Start | End | Strand | Attributes |
|---|---|---|---|---|---|---|
| AgamP4_3L | VEuPathDB | protein_coding_gene | 2248735 | 2251022 | + | ID=AGAP010365;description=unspecified product;ebi_biotype=protein_coding |
| AgamP4_3L | VEuPathDB | protein_coding_gene | 2251295 | 2257902 | - | ID=AGAP010366;description=unspecified product;ebi_biotype=protein_coding |
| AgamP4_3L | VEuPathDB | protein_coding_gene | 2272134 | 2274785 | - | ID=AGAP010368;description=2-hydroxyacyl-CoA lyase 1;ebi_biotype=protein_coding |
| AgamP4_3L | VEuPathDB | protein_coding_gene | 2279614 | 2280171 | - | ID=AGAP010369;Name=CPR112;description=cuticular protein RR-2 family 112;ebi_biotype=protein_coding |
| AgamP4_3L | VEuPathDB | protein_coding_gene | 2277000 | 2285568 | + | ID=AGAP010370;description=solute carrier family 17 (anion/sugar transporter), member 5;ebi_biotype=protein_coding |
| AgamP4_3L | VEuPathDB | protein_coding_gene | 2287747 | 2290635 | - | ID=AGAP010371;description=Glyco_18 domain-containing protein [Source:UniProtKB/TrEMBL;Acc:A0A1S4H4R6];ebi_biotype=protein_coding |
| AgamP4_3L | VEuPathDB | protein_coding_gene | 2301390 | 2318497 | + | ID=AGAP010372;description=solute carrier family 17 (anion/sugar transporter), member 5;ebi_biotype=protein_coding |
| AgamP4_3L | VEuPathDB | protein_coding_gene | 2318591 | 2323312 | - | ID=AGAP010373;description=arginine-tRNA-protein transferase;ebi_biotype=protein_coding |
| AgamP4_3L | VEuPathDB | protein_coding_gene | 2343718 | 2347311 | - | ID=AGAP010376;description=unspecified product;ebi_biotype=protein_coding |
| AgamP4_3L | VEuPathDB | protein_coding_gene | 2369330 | 2371785 | - | ID=AGAP010377;description=unspecified product;ebi_biotype=protein_coding |
| AgamP4_3L | VEuPathDB | protein_coding_gene | 2350719 | 2369594 | + | ID=AGAP010378;description=unspecified product;ebi_biotype=protein_coding |
| AgamP4_3L | VEuPathDB | protein_coding_gene | 2372247 | 2387390 | - | ID=AGAP010379;description=unspecified product;ebi_biotype=protein_coding |
| AgamP4_3L | VEuPathDB | protein_coding_gene | 2388290 | 2392232 | - | ID=AGAP010380;description=CRC domain-containing protein [Source:UniProtKB/TrEMBL;Acc:A0A1S4H4S6];ebi_biotype=protein_coding |
| AgamP4_3L | VEuPathDB | protein_coding_gene | 2393423 | 2395835 | + | ID=AGAP010381;description=unspecified product;ebi_biotype=protein_coding |
| AgamP4_3L | VEuPathDB | protein_coding_gene | 2395885 | 2402310 | - | ID=AGAP010382;Name=Ars2;description=Serrate RNA effector molecule homolog;ebi_biotype=protein_coding |
| AgamP4_3L | VEuPathDB | protein_coding_gene | 2416006 | 2420715 | + | ID=AGAP010383;description=solute carrier family 15 member 1;ebi_biotype=protein_coding |
| AgamP4_3L | VEuPathDB | protein_coding_gene | 2413428 | 2423079 | + | ID=AGAP010384;description=unspecified product;ebi_biotype=protein_coding |
| AgamP4_3L | VEuPathDB | protein_coding_gene | 2430802 | 2433934 | - | ID=AGAP010385;description=unspecified product;ebi_biotype=protein_coding |
| AgamP4_3L | VEuPathDB | protein_coding_gene | 2436355 | 2438695 | - | ID=AGAP010386;description=Translocating chain-associated membrane protein 2;ebi_biotype=protein_coding |
| AgamP4_3L | VEuPathDB | protein_coding_gene | 2444095 | 2447852 | - | ID=AGAP010387;Name=HKT;description=alanine-glyoxylate aminotransferase;ebi_biotype=protein_coding |
| AgamP4_3L | VEuPathDB | protein_coding_gene | 2452624 | 2463174 | - | ID=AGAP010388;description=glucuronyl/N-acetylglucosaminyl transferase EXT1;ebi_biotype=protein_coding |
| AgamP4_3L | VEuPathDB | protein_coding_gene | 2467694 | 2476875 | - | ID=AGAP010389;description=Prestin;ebi_biotype=protein_coding |
| AgamP4_3L | VEuPathDB | protein_coding_gene | 2482177 | 2486705 | - | ID=AGAP010390;Name=COE12O;description=carboxylesterase;ebi_biotype=protein_coding |
| AgamP4_3L | VEuPathDB | protein_coding_gene | 4860623 | 4883826 | - | ID=AGAP010486;Name=GPRALS3;description=Putative allatostatin receptor 3 [Source:UniProtKB/TrEMBL;Acc:A0A1S4H448];ebi_biotype=protein_coding |
| AgamP4_3L | VEuPathDB | protein_coding_gene | 4899887 | 4901343 | + | ID=AGAP010487;description=unspecified product;ebi_biotype=protein_coding |
| AgamP4_3L | VEuPathDB | protein_coding_gene | 4938735 | 4940046 | - | ID=AGAP010488;description=ubiquinone biosynthesis methyltransferase;ebi_biotype=protein_coding |
| AgamP4_3L | VEuPathDB | protein_coding_gene | 4997901 | 4998953 | + | ID=AGAP010489;Name=OBP4;description=odorant-binding protein antennal 4;ebi_biotype=protein_coding |
| AgamP4_3L | VEuPathDB | protein_coding_gene | 5008950 | 5027453 | + | ID=AGAP010490;description=rabconnectin;ebi_biotype=protein_coding |
| AgamP4_3L | VEuPathDB | protein_coding_gene | 5028087 | 5031692 | - | ID=AGAP010491;description=N-acetyltransferase 10;ebi_biotype=protein_coding |
| AgamP4_3L | VEuPathDB | protein_coding_gene | 5031838 | 5034540 | + | ID=AGAP010492;description=asparagine synthetase domain-containing protein c;ebi_biotype=protein_coding |
| AgamP4_3L | VEuPathDB | protein_coding_gene | 5034580 | 5036970 | + | ID=AGAP010493;description=transcription initiation factor TFIID subunit 7;ebi_biotype=protein_coding |
| AgamP4_3L | VEuPathDB | protein_coding_gene | 5037047 | 5041194 | - | ID=AGAP010494;description=partitioning defective protein 6;ebi_biotype=protein_coding |
| AgamP4_3L | VEuPathDB | protein_coding_gene | 5043346 | 5047739 | + | ID=AGAP010496;description=splicing factor, arginine/serine-rich 1/9;ebi_biotype=protein_coding |
| AgamP4_3L | VEuPathDB | protein_coding_gene | 5050649 | 5052019 | + | ID=AGAP010497;Name=Pex12;description=peroxin 12;ebi_biotype=protein_coding |
| AgamP4_3L | VEuPathDB | protein_coding_gene | 5052342 | 5053759 | + | ID=AGAP010498;description=RNA 3'-terminal phosphate cyclase-like protein;ebi_biotype=protein_coding |
| AgamP4_3L | VEuPathDB | protein_coding_gene | 5054214 | 5055709 | + | ID=AGAP010499;Name=AD20590;description=S-(hydroxymethyl)glutathione dehydrogenase [Source:UniProtKB/TrEMBL;Acc:A0A1S4H465];ebi_biotype=protein_coding |

| Sequence | Source | Feature | Start | End | Strand | Attributes |
|---|---|---|---|---|---|---|
| AgamP4_3L | VEuPathDB | protein_coding_gene | 5055968 | 5058317 | - | ID=AGAP010500;description=unspecified product;ebi_biotype=protein_coding |
| AgamP4_3L | VEuPathDB | protein_coding_gene | 5058906 | 5060839 | + | ID=AGAP010501;description=unspecified product;ebi_biotype=protein_coding |
| AgamP4_3L | VEuPathDB | protein_coding_gene | 5064690 | 5065514 | - | ID=AGAP010502;description=unspecified product;ebi_biotype=protein_coding |
| AgamP4_3L | VEuPathDB | protein_coding_gene | 5066089 | 5118810 | - | ID=AGAP010503;description=potassium intermediate/small conductance calcium-activated channel subfamily N;ebi_biotype=protein_coding |
| AgamP4_3L | VEuPathDB | protein_coding_gene | 5124949 | 5126370 | - | ID=AGAP010504;Name=Or43;description=odorant receptor 43;ebi_biotype=protein_coding |
| AgamP4_3L | VEuPathDB | protein_coding_gene | 5133258 | 5134698 | - | ID=AGAP010505;Name=Or44;description=odorant receptor 44;ebi_biotype=protein_coding |
| AgamP4_3L | VEuPathDB | protein_coding_gene | 5182242 | 5212325 | - | ID=AGAP010506;description=unspecified product;ebi_biotype=protein_coding |
| AgamP4_3L | VEuPathDB | protein_coding_gene | 5238515 | 5240204 | + | ID=AGAP010507;description=unspecified product;ebi_biotype=protein_coding |
| AgamP4_3L | VEuPathDB | protein_coding_gene | 5241964 | 5243831 | + | ID=AGAP010508;description=proton-coupled amino acid transporter;ebi_biotype=protein_coding |
| AgamP4_3L | VEuPathDB | protein_coding_gene | 5243916 | 5251412 | - | ID=AGAP010509;description=organic cation transporter 3;ebi_biotype=protein_coding |
| AgamP4_3L | VEuPathDB | protein_coding_gene | 5260848 | 5284114 | + | ID=AGAP010510;description=Tubulin beta chain [Source:UniProtKB/TrEMBL;Acc:Q7PPR6];ebi_biotype=protein_coding |
| AgamP4_3L | VEuPathDB | protein_coding_gene | 5451009 | 5529279 | - | ID=AGAP010513;description=putative muscarinic acetylcholine receptor 1;ebi_biotype=protein_coding |
| AgamP4_3L | VEuPathDB | protein_coding_gene | 5546436 | 5562155 | - | ID=AGAP010514;description=activator of 90 kDa heat shock protein ATPase;ebi_biotype=protein_coding |
| AgamP4_3L | VEuPathDB | protein_coding_gene | 5567345 | 5568203 | - | ID=AGAP010515;description=pre-mRNA-splicing factor SYF2;ebi_biotype=protein_coding |
| AgamP4_3L | VEuPathDB | protein_coding_gene | 5568814 | 5581540 | + | ID=AGAP010516;description=unspecified product;ebi_biotype=protein_coding |
| AgamP4_3L | VEuPathDB | protein_coding_gene | 5581693 | 5583118 | - | ID=AGAP010517;Name=MnSOD1;description=manganese-iron superoxide dismutase 1;ebi_biotype=protein_coding |
| AgamP4_3L | VEuPathDB | protein_coding_gene | 5583730 | 5587129 | - | ID=AGAP010518;description=unspecified product;ebi_biotype=protein_coding |
| AgamP4_3L | VEuPathDB | protein_coding_gene | 5610334 | 5633490 | + | ID=AGAP010519;description=kinesin-like protein unc-104;ebi_biotype=protein_coding |
| AgamP4_3L | VEuPathDB | protein_coding_gene | 5636019 | 5640100 | - | ID=AGAP010520;description=krueppel-like factor, other;ebi_biotype=protein_coding |
| AgamP4_3L | VEuPathDB | protein_coding_gene | 5676660 | 5680124 | + | ID=AGAP010530;Name=CLIPE4;description=CLIP-domain serine protease [Source:UniProtKB/TrEMBL;Acc:A0A1S4H4A2];ebi_biotype=protein_coding |
| AgamP4_3L | VEuPathDB | protein_coding_gene | 5714410 | 5715624 | - | ID=AGAP010531;description=fibrinogen-related protein 7;ebi_biotype=protein_coding |
| AgamP4_3R | VEuPathDB | protein_coding_gene | 49143471 | 49143926 | + | ID=AGAP013367;Name=CPR155;description=cuticular protein RR-2 family 155;ebi_biotype=protein_coding |
| AgamP4_3R | VEuPathDB | protein_coding_gene | 49116463 | 49117244 | + | ID=AGAP013723;description=unspecified product;ebi_biotype=protein_coding |
| AgamP4_3R | VEuPathDB | protein_coding_gene | 49219789 | 49220479 | + | ID=AGAP013749;description=unspecified product;ebi_biotype=protein_coding |
| AgamP4_3L | VEuPathDB | protein_coding_gene | 1333415 | 1391875 | - | ID=AGAP027982;description=Tetraspanin [Source:UniProtKB/TrEMBL;Acc:A0A1S4HDP2];ebi_biotype=protein_coding |
| AgamP4_3L | VEuPathDB | protein_coding_gene | 844271 | 850519 | - | ID=AGAP027999;description=aquaporin;ebi_biotype=protein_coding |
| AgamP4_3L | VEuPathDB | protein_coding_gene | 1424181 | 1429215 | - | ID=AGAP028083;description=DOMON domain-containing protein [Source:UniProtKB/TrEMBL;Acc:A0A1S4HCS6];ebi_biotype=protein_coding |
| AgamP4_3L | VEuPathDB | protein_coding_gene | 2351022 | 2354300 | - | ID=AGAP028146;description=BTB domain-containing protein [Source:UniProtKB/TrEMBL;Acc:A0A1S4HCY8];ebi_biotype=protein_coding |
| AgamP4_3L | VEuPathDB | protein_coding_gene | 4903117 | 4921104 | - | ID=AGAP028205;description=unspecified product;ebi_biotype=protein_coding |
| AgamP4_3L | VEuPathDB | protein_coding_gene | 2448546 | 2449463 | - | ID=AGAP028223;description=unspecified product;ebi_biotype=protein_coding |
| AgamP4_3L | VEuPathDB | protein_coding_gene | 837955 | 843075 | - | ID=AGAP028491;Name=AQP2;description=aquaporin;ebi_biotype=protein_coding |
| AgamP4_3L | VEuPathDB | protein_coding_gene | 1052157 | 1053313 | + | ID=AGAP028493;description=unspecified product;ebi_biotype=protein_coding |
| AgamP4_3L | VEuPathDB | protein_coding_gene | 2435439 | 2436427 | + | ID=AGAP028494;description=unspecified product;ebi_biotype=protein_coding |
| AgamP4_3L | VEuPathDB | protein_coding_gene | 5420400 | 5436061 | + | ID=AGAP028497;description=unspecified product;ebi_biotype=protein_coding |
| AgamP4_3L | VEuPathDB | protein_coding_gene | 5438584 | 5452753 | + | ID=AGAP028498;description=unspecified product;ebi_biotype=protein_coding |
| AgamP4_3L | VEuPathDB | protein_coding_gene | 1266789 | 1302384 | + | ID=AGAP028612;description=unspecified product;ebi_biotype=protein_coding |
| AgamP4_3L | VEuPathD | protein_coding_g | 2341035 | 2341997 | + | ID=AGAP028693;Name=RpS18;description=40S ribosomal protein |

| Sequence | Source | Feature | Start | End | Strand | Attributes |
|---|---|---|---|---|---|---|
| | B | ene | | | | S18;ebi_biotype=protein_coding |
| AgamP4_3L | VEuPathDB | protein_coding_gene | 2324923 | 2325497 | + | ID=AGAP028694;Name=RpS18;description=40S ribosomal protein S18;ebi_biotype=protein_coding |
| AgamP4_3R | VEuPathDB | protein_coding_gene | 49146965 | 49147426 | + | ID=AGAP029247;Name=CPR148;description=cuticular protein RR-2 family 148;ebi_biotype=protein_coding |
| AgamP4_3R | VEuPathDB | protein_coding_gene | 49145656 | 49146108 | + | ID=AGAP029248;Name=CPR156;description=cuticular protein RR-2 family 156;ebi_biotype=protein_coding |
| AgamP4_3L | VEuPathDB | protein_coding_gene | 2233385 | 2233834 | - | ID=AGAP029251;description=Chitin-binding type-2 domain-containing protein [Source:UniProtKB/TrEMBL;Acc:A0A453YZ26];ebi_biotype=protein_coding |
| AgamP4_3L | VEuPathDB | protein_coding_gene | 2225850 | 2226986 | - | ID=AGAP029275;description=unspecified product;ebi_biotype=protein_coding |
| AgamP4_3L | VEuPathDB | protein_coding_gene | 2507869 | 2511896 | + | ID=AGAP029278;description=NACHT domain-containing protein [Source:UniProtKB/TrEMBL;Acc:A0A453YZJ4];ebi_biotype=protein_coding |
| AgamP4_3L | VEuPathDB | protein_coding_gene | 2223788 | 2225488 | - | ID=AGAP029283;description=unspecified product;ebi_biotype=protein_coding |
| AgamP4_3L | VEuPathDB | protein_coding_gene | 2357118 | 2358069 | - | ID=AGAP029291;description=unspecified product;ebi_biotype=protein_coding |
| AgamP4_3L | VEuPathDB | protein_coding_gene | 4995203 | 5000405 | + | ID=AGAP029303;description=unspecified product;ebi_biotype=protein_coding |
| AgamP4_3R | VEuPathDB | protein_coding_gene | 52400373 | 52401100 | + | ID=AGAP029400;description=6PGD domain-containing protein [Source:UniProtKB/TrEMBL;Acc:A0A453YZI6];ebi_biotype=protein_coding |
| AgamP4_3L | VEuPathDB | protein_coding_gene | 5313190 | 5342479 | + | ID=AGAP029625;description=MADS-box domain-containing protein [Source:UniProtKB/TrEMBL;Acc:A0A453Z090];ebi_biotype=protein_coding |
| AgamP4_3L | VEuPathDB | protein_coding_gene | 2124999 | 2140343 | + | ID=AGAP029666;description=RING-type domain-containing protein [Source:UniProtKB/TrEMBL;Acc:A0A453Z0L3];ebi_biotype=protein_coding |
| AgamP4_3L | VEuPathDB | protein_coding_gene | 2232300 | 2232629 | - | ID=AGAP029714;description=Chitin-binding type-2 domain-containing protein [Source:UniProtKB/TrEMBL;Acc:A0A453Z0J3];ebi_biotype=protein_coding |
| AgamP4_3L | VEuPathDB | protein_coding_gene | 2235020 | 2236107 | - | ID=AGAP029715;description=unspecified product;ebi_biotype=protein_coding |
| AgamP4_3L | VEuPathDB | protein_coding_gene | 2264147 | 2265016 | + | ID=AGAP029717;description=unspecified product;ebi_biotype=protein_coding |
| AgamP4_3R | VEuPathDB | protein_coding_gene | 52152185 | 52152607 | + | ID=AGAP029880;description=unspecified product;ebi_biotype=protein_coding |
| AgamP4_3L | VEuPathDB | protein_coding_gene | 901312 | 1157072 | - | ID=AGAP029957;description=unspecified product;ebi_biotype=protein_coding |
| AgamP4_3L | VEuPathDB | protein_coding_gene | 1311377 | 1311999 | - | ID=AGAP029958;description=unspecified product;ebi_biotype=protein_coding |
| AgamP4_3L | VEuPathDB | protein_coding_gene | 1673430 | 1675835 | + | ID=AGAP029959;description=unspecified product;ebi_biotype=protein_coding |
| AgamP4_3L | VEuPathDB | protein_coding_gene | 2267287 | 2272039 | + | ID=AGAP029980;description=unspecified product;ebi_biotype=protein_coding |

### Table S19. Genome assemblies and annotation resources used in this study

| ***Anopheles epiroticus*** | **https://vectorbase.org/vectorbase/app/downloads/Current_Release/AepiroticusEpiroticus2/** |
|---|---|
| *Anopheles christyi* | https://vectorbase.org/vectorbase/app/downloads/Current_Release/AchristyiACHKN1017/ |
| *Anopheles arabiensis* | https://vectorbase.org/vectorbase/app/downloads/Current_Release/AarabiensisDONGOLA |
| *Anopheles coluzzii* | https://vectorbase.org/vectorbase/app/downloads/Current_Release/AcoluzziiAcolN3/ |
| *Anopheles gambiae* | https://vectorbase.org/vectorbase/app/downloads/Current_Release/AgambiaePEST |
| *Anopheles melas* | https://vectorbase.org/vectorbase/app/downloads/Current_Release/AmelasCM1001059_A/ |
| *Anopheles merus* | https://vectorbase.org/vectorbase/app/downloads/Current_Release/AmerusMAF |
| *Anopheles quadriannulatus* | https://vectorbase.org/vectorbase/app/downloads/Current_Release/AquadriannulatusSANGWE/ |

**Table S20. Sample IDs of the Ag1000G phase 3 individuals used in population genomic analyses. Each species is represented by 59 individuals**

| *A. gambiae* | *A. coluzzii* | *A. arabiensis* |
|---|---|---|
| AB0373-C | AB0326-C | AB0333-C |
| AB0374-C | AB0327-C | AB0457-C |
| AB0375-C | AB0328-C | AB0502-C |
| AB0378-C | AB0329-C | AN0341-C |
| AB0379-C | AB0330-C | AN0338-C |
| AB0381-C | AB0331-C | AK0041-C |
| AB0386-C | AB0332-C | AK0042-C |
| AB0388-C | AB0334-C | AK0043-C |
| AB0389-C | AB0337-C | AK0044-C |
| AB0432-C | AB0338-C | AK0046-C |
| AB0433-C | AB0339-C | AK0047-C |
| AB0435-C | AB0340-C | AK0054-C |
| AB0436-C | AB0341-C | AK0055-C |
| AB0439-C | AB0348-C | AK0056-C |
| AB0440-C | AB0351-C | AK0057-Cx |
| AB0458-C | AB0352-C | AK0284-C |
| AB0459-C | AB0354-C | AK0298-C |
| AB0460-C | AB0355-C | AK0299-C |
| AB0462-C | AB0356-C | AZ0152-C |
| AB0463-C | AB0357-C | AZ0153-C |
| AB0465-C | AB0358-C | AZ0155-C |
| AB0466-C | AB0359-C | AZ0156-C |
| AB0467-C | AB0360-C | AZ0157-C |
| AB0468-C | AB0361-C | AZ0158-C |
| AB0469-C | AB0362-C | AZ0159-C |
| AB0470-C | AB0363-C | AZ0160-C |
| AB0471-C | AB0364-C | AZ0161-C |
| AB0472-C | AB0365-C | AZ0164-C |
| AB0473-C | AB0366-C | AZ0165-C |
| AB0474-C | AB0367-C | AZ0166-C |
| AB0477-C | AB0368-C | AZ0167-C |
| AB0504-C | AB0369-C | AZ0168-C |
| AB0517-C | AB0370-C | AZ0170-C |
| AB0524-C | AB0371-C | AZ0172-C |
| AB0525-C | AB0372-C | AZ0174-C |
| AB0526-C | AB0396-C | AZ0175-C |
| AB0527-C | AB0397-C | AZ0176-C |
| AB0528-C | AB0398-C | AZ0179-C |
| AB0529-C | AB0399-C | AZ0180-C |
| AB0530-C | AB0400-C | AZ0181-C |
| AB0531-C | AB0403-C | AZ0182-C |
| AB0532-C | AB0404-C | AZ0183-C |
| AB0533-C | AB0409-C | AZ0184-C |
| AB0536-C | AB0410-C | AZ0185-C |
| AB0537-C | AB0411-C | AZ0186-C |
| AB0538-C | AB0413-C | AZ0187-C |
| AN0325-C | AB0503-C | AZ0188-C |
| AN0408-C | AB0505-C | AZ0191-C |
| AN0326-C | AB0508-C | AZ0192-C |
| AN0585-CW | AB0509-C | AZ0199-C |
| AN0577-CW | AB0510-C | AZ0200-C |
| AN0586-CW | AB0523-C | AZ0201-C |
| AN0578-CW | AB0408-C | AZ0202-C |
| AN0587-CW | AN0348-C | AZ0223-C |
| AN0579-CW | AN0402-C | AZ0226-C |
| AN0588-CW | AN0344-C | AZ0242-C |

| *A. gambiae* | *A. coluzzii* | *A. arabiensis* |
|---|---|---|
| AN0580-CW | AN0379-C | AZ0246-C |
| AN0581-CW | AN0332-C | AZ0247-C |
| AN0574-CW | AN0601-CW | AZ0251-C |
